# Plexin-B1 mutation drives prostate cancer metastasis

**DOI:** 10.1101/2020.07.19.203695

**Authors:** Boris Shorning, Neil Trent, David Griffiths, Thomas Worzfeld, Stefan Offermanns, John Masters, Alan R. Clarke, Matthew J. Smalley, Magali Williamson

## Abstract

Prostate cancer mortality is associated with the metastatic spread of tumour cells. A better understanding of the mechanisms which allow a locally advanced tumour to disseminate around the body will identify new therapeutic targets to block this process. One of set of genes implicated in metastasis are plexins, which can promote or suppress tumour progression depending on cancer type and cellular context. We have taken a mouse genetics approach to gain insight into the role of Plexin-B1 in prostate cancer progression *in vivo*.

We show here that genetic deletion of Plexin-B1 in *PbCre^+^Pten^fl/fl^Kras^G12V^* and *PbCre^+^Pten^fl/fl^p53^fl/fl^* mouse prostate cancer models significantly decreased metastasis. High levels of prostate epithelial cell-specific expression of wild-type Plexin-B1 in knock-in mice with a *PbCre^+^Pten^fl/fl^Kras^G12V^* background also significantly decreased metastasis. In contrast, expression of a Plexin-B1 mutant (P1597L; identified from metastatic deposits in prostate cancer patients) in prostate epithelial cells in *PbCre^+^Pten^fl/fl^Kras^G12V^* and *PbCre^+^Pten^fl/fl^p53^fl/fl^* mice significantly increased metastasis, in particular metastasis to distant sites. In line with these findings, both deletion and overexpression of wild-type Plexin-B1 reduced invasion of tumour cells into the prostate stroma, while overexpression of mutant Plexin-B1 significantly increased invasion, suggesting that Plexin-B1 has a role in the initial stages of metastasis. Invasion and metastasis also correlated with phosphorylation of myosin light chain, suggesting that Plexin-B1 signals via the Rho/ROCK pathway to promote metastasis.

Our results demonstrate that mutant Plexin-B1 promotes metastasis in prostate cancer and represents a new therapeutic target to suppress tumour spread.

## Introduction

Metastasis is the primary cause of morbidity and mortality in prostate cancer, yet despite increasing interest in treating oligometastatic disease with curative intent(1), few effective treatments specifically developed to counter metastasis are currently available in the clinic. It is also not clear what drives prostate cancer progression from locally advanced to invasive/disseminated disease but it is likely to result from the activation of a complex combination of alternative signalling pathways(2). Studies using *in vivo* models of prostate cancer of different genetic backgrounds allow the role of individual pathways contributing to the process to be defined and potential therapeutic targets identified.

The early stages of metastasis involve the migration and invasion of tumour cells out of the primary tumour through the basement membrane and beyond. One of set of genes implicated in this process are plexins, cell surface receptors for semaphorins(3). Vertebrates possess nine plexin genes, classified into 4 subfamilies (class A(1-4), B(1-3) C1 and D1)(3). Plexin stimulation delivers directional cues for cell migration and axon guidance through the regulation of several small GTPases and semaphorin-plexin signalling can either be attractive or repulsive depending on the particular plexin co-receptor expressed(4, 5), while non-polarised stimulation of cells with semaphorins results in cell collapse(6). B-class plexins can interact with the GTPases Rnd1-3(7, 8), Rac(9), RhoD(10), R-Ras(7) and M-Ras(11) and regulate Rho via PDZRhoGEF/LARG(12) and p190RhoGAP(13). In addition, the plexin cytoplasmic tail contains a GTPase activating protein (GAP) domain(14) divided into two regions by a Rho Binding domain (RBD)(15) and plexins act as GAPs for Rap1B and Rap2A(16). PlexinB1 is a tumour suppressor gene in some types of cancer (e.g ER-positive breast cancer)(17) but promotes tumour progression in other types (e.g. ERBB2-expressing breast cancer)(18) while somatic missense mutations in *PLXNB1* occur in patient samples of prostate cancer metastasis(19). Functional analysis *in vitro* of three such mutations (T1697A, T1795A, L1815P), demonstrated that these sequence changes inhibit the interaction of Rnd1, Rac and R-Ras with Plexin-B1 and block the ability of Plexin-B1 to mediate R-Ras inactivation(19, 20). Overexpression of these three mutations, along with another GAP domain mutation (P1597L) **(Supplementary Figure 1A)**, increased cell motility, invasion and anchorage-independent growth of prostate cancer cells *in vitro*. In contrast overexpression of wild-type Plexin-B1 reduced cell motility and invasion(19). A better understanding of the role of mutant Plexin-B1 in metastatic prostate cancer, and the downstream pathways it activates, offers the potential for novel therapeutic approaches in this setting.

To gain *in vivo* insights into the cellular mechanisms involved in metastasis and to investigate the role of Plexin-B1 in this process, we genetically manipulated the expression of wild-type and mutant Plexin-B1 in two models of prostate cancer and studied the effect on prostate cancer progression *in vivo*. We find that Plexin-B1 plays a significant role in the spread of prostate cancer, with high expression of the wild type protein suppressing metastasis but expression of the clinically relevant P1597L mutant enhancing it, likely through activation of the Rho/ROCK pathway. This pathway has potential as a therapeutic target in locally advanced or oligometastatic prostate cancer, particularly when mutant Plexin-B1 is present.

## Methods

### Experimental animals

All animal studies and breeding were carried out under UK Home Office regulations and ARRIVE guidelines for the care and use of animals(21). Research was hypothesis and objective-driven to minimize the number of animals used but power calculations ensured sufficient animals were included in cohorts to gain statistically significant results. *Pb-Cre^+^* (*Pb-Cre4, ARR2PB*) mice were obtained from the Mouse Models of Human Cancers Consortium (National Cancer Institute-Frederick). The *PB-Cre* transgene was incorporated into cohorts using male mice, as *PB-Cre^+^* female mice have been shown to recombine in the ovaries(22). Littermate controls lacking the *Pb-Cre* transgene were used in all experiments. Mice homozygous for floxed *Pten* exons 4 and 5 (*Pten^fl/fl^*)(23), mice carrying inducible endogenous *Kras^G12V^* oncogene(24), mice homozygous for floxed *p53* exons 2-10 (*p53^fl/fl^*)(25) and constitutive *Plexin-B1*–knockout (*PlxnB1^-/-^*) mice lacking exons 13-16(26) have been described previously. Mice were maintained on an outbred background.

### Genetically modified mice

Mice carrying a conditionally activated knock-in construct of human Plexin-B1 containing wild-type or a C5060T mutation causing a substitution of proline 1597 to leucine (*loxP-STOP-loxP-PLXNB1^WT^* or *loxP-STOP-loxP-PLXNB1^P1597L^* mice; hereafter *PLXNB1^WT^* and *PLXNB1^MUT^*) were commercially generated by genOway (Lyon, France). The cDNA construct *P1597L-PLXNB1* had been made and sequenced previously(19). A transgenic cassette expressing the wild-type or mutated *P1597L-PLXNB1 cDNA – hGH polyA* under the control of the *CAG* promoter and with a neomycin STOP sequence flanked by two *loxP* sites was generated. GenOway’s validated *Rosa26* “Quick Knock-in™” approach was then used to introduce a single copy of the cassette into the *Rosa26* locus on chromosome 6 through homologous recombination in embryonic stem cells (**Supplementary Figure 1B-D**). The linearized construct was transfected into mouse 129Sv ES cells according to standard electroporation procedures (i.e. 5×10^6^ ES cells in presence of 40 μg of linearized plasmid, 260 volt, 500 μF). Positive selection was started 48 hours after electroporation by addition of 200 μg/ml of G418. G418-resistant clones were screened for the correct homologous recombination event at the *Rosa26* locus by PCR and southern blotting. Three correctly recombined ES clones were used for injection into C57BL/6J blastocysts. Injected blastocysts were re-implanted into OF1 pseudo-pregnant females and allowed to develop to term. After approximately three weeks a total of 3 male chimeric mice were produced per construct with a chimerism rate above 50%. These animals were mated with WT C57BL/6J females to generate heterozygous mice carrying the *Rosa26* floxed allele in their germline.

To assess whether the ES cells contributed to the germ layer of the chimeras, mouse coat colour markers were used. The coat colour marker of the 129Sv ES cells (agouti) is dominant over the black coat colour of the C57BL/6 mice. Agouti F1 progeny were screened for the *Rosa26* knock-in allele by PCR and southern blotting, using genomic DNA isolated from tail biopsies. 9 animals out of 30 carried the allele and 6 animals were verified further by southern blotting (**Supplementary Figure 1B-D**).

*PLXNB1^WT^* and *PLXNB1^MUT^* knock-in animals were bred to *Pb-Cre^+^* mice to obtain animals with prostate epithelial specific expression of wild-type or mutant PlexinB1 protein. Mice were genotyped for the *PLXNB1^WT^* or *PLXNB1^MUT^*-inducible allele (heterozygous or homozygous) by PCR either according to a genOway protocol (forward *AAGACGAAAAGGGCAAGCATCTTCC*, reverse *GCAGTGAGAAGAGTACCACCATGAGTCC*, 94°C for 2 min and 35 cycles of 94°C for 30”, 65°C for 30”, 68°C for 5 min, giving a 1870 bp product in the inducible *PLXNB1^MUT^* mouse) or according to our simplified protocol identical to quantitative Real-Time PCR analysis of *PlxnB1* expression. To distinguish between heterozygous and homozygous *PLXNB1^MUT^* we followed a protocol developed by genOway (forward *CAATACCTTTCTGGGAGTTCTCTGC*, reverse *CTGCATAAAACCCCAGATGACTACC*, 94°C for 2 min and 35 cycles of 94°C for 30’’, 55°C for 30’’, 72°C for 30’’, giving a 304 bp product in WT or heterozygous *PLXNB1^MUT^* mouse and no product in homozygous *PLXNB1^MUT^* mouse).

### RNA extraction and quantitative Real-Time rtPCR analysis

Prostate lobes were dissected in ice-cold PBS. Tissues were homogenised in TRIZOL Reagent (Invitrogen, Paisley, UK), extracted using standard phenol–chloroform protocol and RNA was purified further using an RNA extraction kit (Norgen, Thorold, Ontario, Canada). RNA from dorsolateral lobes of age-matched 100day old mice (n=2 of each genotype) were used for quantitative Real-Time rtPCR (qRTPCR). Reverse transcription was performed using the SuperScript III reverse transcriptase kit and random hexamers (Invitrogen) according to the manufacturer’s instructions. SYBR Select Master Mix (Applied Biosystems, Fisher Scientific, Loughborough, UK) was added to cDNA samples and primers. Samples were run using QuantStudio 6 Flex Real-Time PCR System (Applied Biosystems). Reverse transcriptase negative controls were included in all analyses. PlexinB1 primers were used for detecting both endogenous *PlxnB1* and *PLXNB1^MUT^* transcripts (forward-*TGTCACTATCAGGGGCTCCA*, reverse-*CTCCCCGCTGGCTCCAGTGAT*, 94°C for 2 min and 35 cycles of 94°C for 30’’, 55°C for 30’’, 72°C for 30’’, giving 145 bp products for both *WT* and inducible *PLXNB1^MUT^*). *β-Actin* and *GAPDH* were used as reference genes.

### Histological analysis and immunohistochemistry (IHC)

Prostate tissue was dissected in 1xPBS and fixed in ice-cold 10% neutral buffered formalin for no longer than 24 h before being processed into paraffin blocks according to standard procedures. For IHC, 5μm sections were dewaxed in xylene, rehydrated in ethanol and antigen retrieval was performed by heating in either citrate (pH 6.0) or EDTA buffer (pH 8.0) in a pressure cooker for 15 min after reaching full pressure. Sections were cooled for 15 min, blocked in 0.5% hydrogen peroxide for 5 minutes at room temperature and then blocked with 20% normal rabbit or goat serum (DAKO, Agilent, Cheadle, Cheshire, UK) for 20 min and incubated with the primary antibody overnight at +4°C. After washing in TBS/0.05% Tween, sections were incubated in secondary antibody for 30 min (EnVision+ System-HRP Labelled Polymer; Dako) and the staining was visualized with DAB (EnVision+ System). Ki67, phospho-MLC2^Ser19^ and androgen receptor (AR) staining were each quantified in prostates of 100day old mice (for each stain n=3 per cohort, 5 fields for each sample). Ki67 staining was scored as positive or negative and compared to the total number of nuclei in a field. For semiquantitative analysis of phospho-MLC2^Ser19^ staining we used a Histo-score (H-score) formula: 3 x percentage of strongly staining cells + 2 x percentage of moderately staining cells + percentage of weakly staining cells, giving a range of 0 to 300.

### Western blotting

Prostate tissues were isolated and homogenized in ice-cold RIPA buffer (150 mM sodium chloride, 1% NP-40, 0.5% sodium deoxycholate, 0.1% SDS, 50 mM Tris, pH 8.0, 1 mM PMSF, 1 μg/ml aprotinin, 1 μg/ml leupeptin). Protein concentrations were measured using a protein assay kit (Pierce™ BCA Protein Assay Kit, #23225, ThermoFisher Scientific, Cheshire, UK). Protein samples were resolved through 10% SDS-polyacrylamide gel and transferred to nitrocellulose membrane. After blocking in 5% non-fat milk, membranes were probed with primary antibodies. Protein detection was performed with ECL reagents according to the manufacturer’s protocol using ChemiDoc XRS System and Image Lab software (Bio-Rad, Watford, Hertfordshire, UK). β-actin was used to normalize sample loading.

### Antibodies

The following antibodies were used in this study: Anti-Plexin-B1 (rabbit polyclonal; Santa Cruz Biotechnology, Heidelberg, Germany; #sc-25642; 1:2000 for IHC; 1:500 for western blot). Anti-pan-cytokeratin AE1/AE3 (mouse monoclonal; Santa Cruz Biotechnology; #sc-81714; 1:250 for IHC). Anti-androgen receptor N-20 (rabbit polyclonal; Santa Cruz Biotechnology; #sc-816; 1:200 for IHC). Anti-phospho-MLC2-Ser19 (rabbit polyclonal; Cell Signalling Technology, Leiden, The Netherlands; #3671; 1:100 for IHC). Anti-β-actin (mouse monoclonal; Sigma-Aldrich, Poole, Dorset, UK; 1:12000 for western blot). Anti-Ki-67 (mouse monoclonal; Cell Signalling Technology; #9449; 1:400 for IHC).

## Results

### Establishing mouse models of PlexinB1 overexpression

To understand the contribution of Plexin-B1 mutation to prostate cancer progression, we established two lines of mice carrying a targeted insertion of either *PLXNB1^WT^* or *PLXNB1^MUT^* cDNA preceded by a *flox-STOP-flox* cassette into the *Rosa26* locus. Targeting was confirmed by Southern blotting (**Supplementary Figure 1**). Activation/over-expression of these conditional alleles in the prostate was achieved by crossing with a line in which CRE recombinase was expressed under the *Probasin* promoter (see **Methods**). Expression of PlexinB1 in prostates of *PLXNB1^WT^* and *PLXNB1^MUT^* mice was compared to unmanipulated wild type mice and to germline homozygous deleted *PlxnB1* mice (*PlxnB1^-/-^*)(26). Plexin-B1 protein was expressed in the epithelial cells of all lobes of wild-type mouse prostates, localising to the cell membrane, cytoplasm and nucleus of prostate epithelial cells, but was absent from prostate stroma **(Figure 1A)**. Plexin-B1 expression was absent from all tissues in *PlxnB1^-/-^* mice, as expected (**Figure 1B**) but high levels of expression of Plexin-B1 were observed in prostate epithelial cells of *PLXNB1^WT^* (**Figure 1C**) and *PLXNB1^MUT^* (**Figure 1D**) mice. The loss of Plexin-B1 expression in *PlxnB1^-/-^* mice, and its overexpression in *PLXNB1^MUT^* prostates, was confirmed by quantitative real time rtPCR gene expression analysis and western blotting **(Figure 1E,F)**. No obvious structural or histological changes in the prostate were found between wild-type and *PlxnB1^-/-^, PLXNB1^WT^* or *PLXNB1^MUT^* mice. Mice from all three lines were fertile and survived over 500 days (as reported previously for *PlxnB1^-/-^* mice).

**Figure 1.**
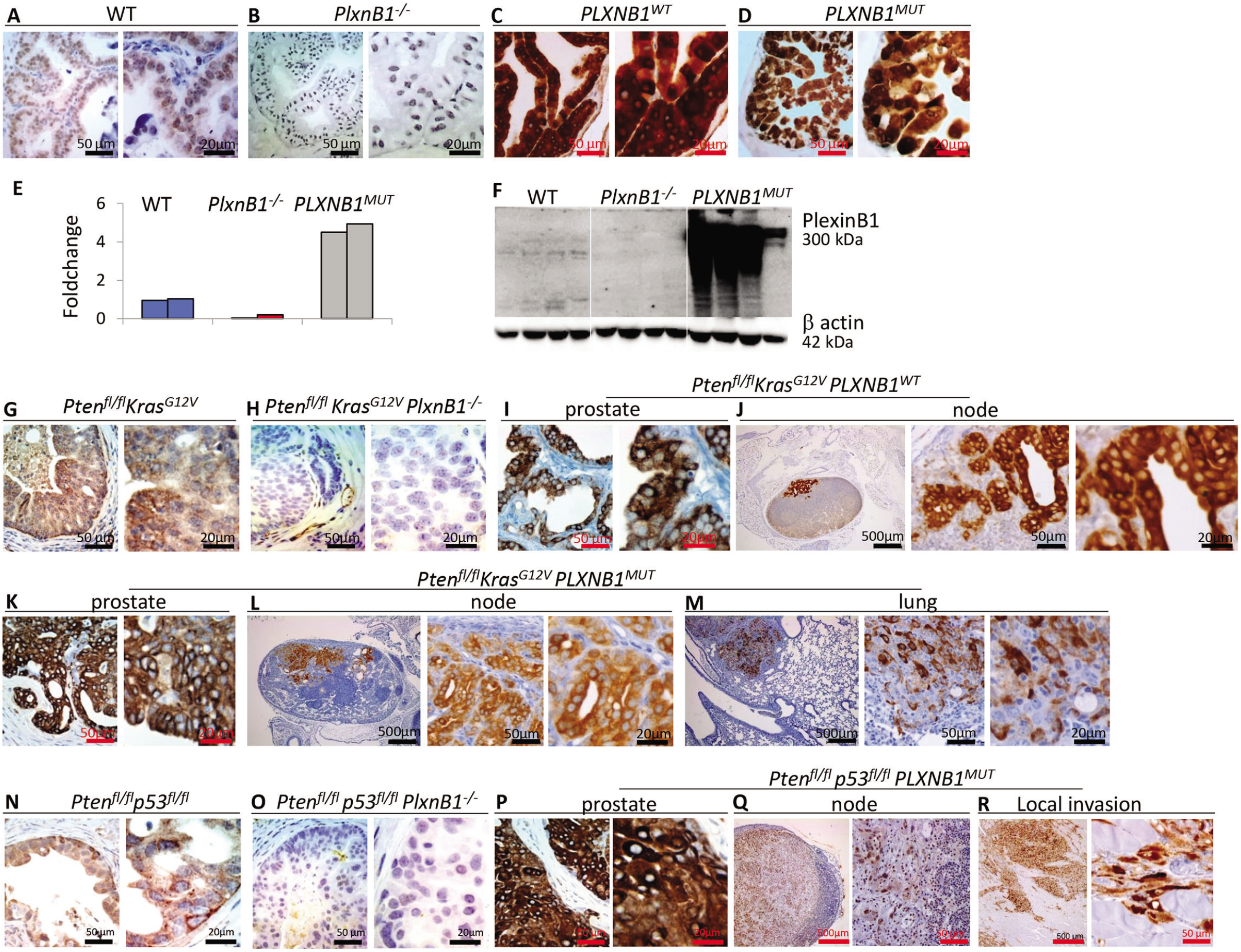
PlexinB1 is expressed in mouse prostate, prostate tumours and metastatic deposits. (**A-D**) Immunostaining for PlexinB1 protein in prostates of WT **(A)**, *PlxnB1^-/-^* **(B)**, *PLXNB1^WT^* **(C)** and *PLXNB1^MUT^***(D)** mice, showing lack of PlexinB1 staining in *PlxnB1^-/-^* mice and high levels of epithelial-specific PlexinB1 expression in prostate of *PLXNB1^WT^* and *PLXNB1 ^P1597L^* mice. **(E)** PlexinB1 mRNA expression in WT, *PlxnB1^-/-^* and *PLXNB1^MUT^* mice (quantitative Real-Time rtPCR of two animals from each genotype, with fold changes compared to an average of the two WT samples). **(F)** Western blot analysis of PlexinB1 protein expression in WT, *PlxnB1^-/-^* and *PLXNB1^MUT^* mouse prostates. Plexin-B1-deficient mice show no PlexinB1 expression; 100 day old mice, 4 animals per genotype. **(G)** PlexinB1 expression in prostate epithelium of *Pten^fl/fl^Kras^G12V^* mice, showing prostate epithelial cell expression. **(H)** Lack of PlexinB1 expression in *Pten^fl/fl^Kras^G12V^PlxnB1^-/-^* mice. (**I, J)**, high levels of epithelial specific PlexinB1 expression in prostate **(I)** and in a lymph node metastasis **(J)** of *Pten^fl/fl^Kras^G12V^PLXNB1^WT^* mice. **(K-M)**, High levels of epithelium-specific PlexinB1 expression in prostate **(K)**, lymph node metastasis **(L)**, and in lung metastasis **(M)** of *Pten^fl/fl^Kras^G12V^PLXNB1^MUT^* mice. **(N)** PlexinB1 expression in prostate epithelium of *Pten^fl/fl^ p53^fl/fl^* mice. **(O)** Lack of PlexinB1 expression in *Pten^fl/fl^p53^fl/fl^PlxnB1^-/-^* mice. **(P-R)** high levels of epithelial specific PlexinB1 expression in prostate **(P),** lymph node metastasis **(Q)** and locally invading cells **(R)** of *Pten^fl/fl^ p53^fl/fl^PLXNB1^MUT^* mice. Scale bars indicated in individual panels.

### PlexinB1^MUT^ overexpression suppresses proliferation in primary mouse prostate tumours but promotes histological heterogeneity in metastatic deposits

Prostate tumours are characterised by loss of PTEN and dysregulation of the PI3K/AKT pathway(27, 28), p53 deletion or mutation(28), and activation of the Ras/Raf pathway(27). To determine whether Plexin-B1 contributes to prostate cancer progression *in vivo*, we tested the effect on tumour growth and metastasis of manipulating Plexin-B1 expression in two different transgenic mouse models of prostate cancer which recapitulate these common genetic alterations and which also metastasise: *PbCre^+^Pten^fl/fl^Kras^G12V^*(29) (hereafter abbreviated to *Pten^fl/fl^Kras^G12V^*) and *PbCre^+^Pten^fl/fl^p53^fl/fl^*(30) (hereafter abbreviated to *Pten^fl/fl^p53^fl/fl^*). Both the *Pten^fl/fl^Kras^G12V^* and *Pten^fl/fl^p53^fl/fl^* models have moderate/low metastatic ability that enables the analysis of additional alleles which may accelerate tumour formation/progression, prostate tumours in both models can metastasize to lumbar lymph nodes with limited ability to form distant metastases(29, 30). *Pten^fl/fl^Kras^G12V^* differs from the previously described highly metastatic *PbCre^+^Pten^fl/fl^Kras^G12D^* model(31) in using a less aggressive *Kras^G12V^*(24). The *Pten^fl/fl^Kras^G12V^* line was crossed with *PlxnB1^-/-^* mice, *PLXNB1^WT^* and *PLXNB1^P1597L^* mice and the *Pten^fl/fl^p53^fl/fl^* line was crossed with the *PlxnB1^-/-^* mice and *PLXNB1^P1597L^* mice. Two cohorts were established for each cross, one for euthanasia at a fixed timepoint of 100 days and one for euthanasia when required for welfare reasons due to tumour morbidity. Primary prostate tumours, local lymph nodes and visceral organs were all processed for histological analysis to assess primary tumour morphology, extent of local invasion and the presence of local or distant metastases.

Plexin-B1 was expressed in the epithelial cells of *Pten^fl/fl^Kras^G12V^* prostate tumour cells (**Figure 1G**), absent from *Pten^fl/fl^Kras^G12V^PlxnB1^-/-^* tissue (**Figure 1H**) and expressed at high levels in *Pten^fl/fl^Kras^G12V^PLXNB1^WT^* and *Pten^fl/fl^Kras^G12V^PLXNB1^MUT^* primary tumours and metastases (**Figure 1I-M**). Similarly, PlexinB1 was expressed in *Pten^fl/fl^p53^fl/fl^* tumour cells, not in *Pten^fl/fl^p53^fl/fl^PlxnB1^-/-^* tissue but at high levels in *Pten^fl/fl^p53^fl/fl^PLXNB1^MUT^* primary tumours and metastases (**Figure 1N-R**).

As described previously (Jefferies et al, 2017, Martin et al, 2011) prostates from both *Pten^fl/fl^Kras^G12V^* and *Pten^fl/fl^p53^fl/fl^*-based cohorts displayed distorted glandular structure with focal areas of microinvasion adjacent to reactive stroma and regions of sarcomatoid metaplasia (**Figure 2A-C, 3A-D**). Prostate tumours from all *Pten^fl/fl^p53^fl/fl^*-based cohorts had a marked increase in mesenchymal phenotype with little epithelial component compared to that of *Pten^fl/fl^Kras^G12V^*-based cohorts (**Figure 2A-C, 3A-D**) and these sarcomatoid tumours were the cause of morbidity in accordance with earlier data(30). Although there were no overt differences in histology of primary tumours between the *Pten^fl/fl^Kras^G12V^*, *Pten^fl/fl^Kras^G12V^PlxnB1^-/-^*, *Pten^fl/fl^Kras^G12V^PLXNB1^WT^* and *Pten^fl/fl^Kras^G12V^PLXNB1^MUT^* models (**Figure 2**), the metastatic deposit composition varied between adenocarcinoma, sarcomatoid (low pan-cytokeratin staining) and squamous areas (high pan-cytokeratin staining) **(Supplementary Figures 2-4)**. Adenocarcinoma was the predominant tissue type in *Pten^fl/fl^Kras^G12V^* cohort metastases **(Figure 2J, Supplementary Figure 2)** with one animal with a lymph node metastatic deposit of predominantly squamous differentiation at day 186 (Supplementary Figure 3, B) and one animal with sarcomatoid deposits in nodes, peritoneum and lung at day 246 (**Figure 2K,L; Supplementary Figure 2F**). Metastases in *Pten^fl/fl^Kras^G12V^PlxnB1^-/-^* and *Pten^fl/fl^Kras^G12V^PLXNB1^WT^* mice were adenocarcinomas (**Figure 2 M-O; Supplementary Figure 3**). Metastases in *Pten^fl/fl^Kras^G12V^PLXNB1^MUT^* mice showed a greater heterogeneity (**Figure 2P-V; Supplementary Figure 4**); 6 out 10 *Pten^fl/fl^Kras^G12V^PLXNB1^MUT^* mice that were taken by day 200 developed predominantly sarcomatoid metastases (**Figure 2Q, S,U; Supplementary Figure 4 A-C,E,F,H**) and *Pten^fl/fl^Kras^G12V^PLXNB1^MUT^* mice taken after day 200 displayed mixed adenocarcinoma/squamous differentiation (**Figure 2, Supplementary Figure 4**).

**Figure 2.**
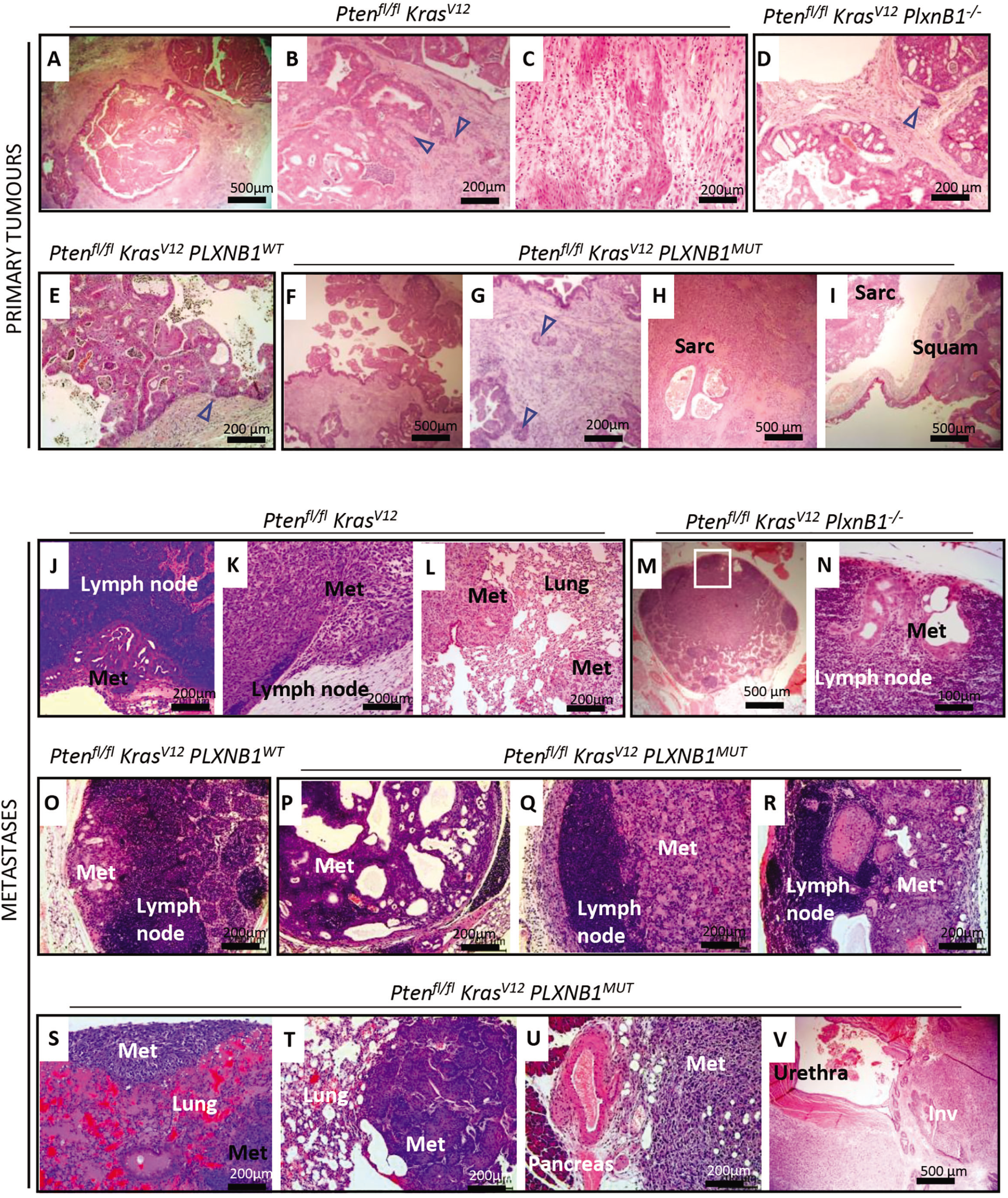
H&E histology of primary prostate tumours and metastatic deposits in *Pten^fl/fl^Kras^G12V^* cohorts (see also Supplementary Figures 3-6). **(A,B)** Invasive adenocarcinoma in *Pten^fl/fl^ Kras^G12V^* prostates at day 100 timepoint before the onset of metastasis; invasive tumour tissue is marked with triangular arrows. **(C)** Aged *Pten^fl/fl^ Kras^G12V^* prostate (6-7 months) showing region of sarcomatoid carcinoma. **(D)** *Pten^fl/fl^Kras^G12V^PlxnB1^-/-^* prostate. **(E)** *Pten^fl/fl^ Kras^G12V^PLXNB1^WT^*. Signs of invasion were rare (marked with an arrow) in these cohorts and stromal reaction was diminished. **(F, G)** Widespread invasion of glandular neoplastic cells (marked with arrows) into the stroma in *Pten^fl/fl^ Kras^G12V^PLXNB1^MUT^* prostates at the day 100 timepoint. **(H,I)** Heterogeneous prostate tumours in aged (>6 months old) *Pten^fl/fl^Kras^G12V^PLXNB1^MUT^* mouse showing sarcomatoid phenotype (**H**) or a mixture of adenomatous, sarcomatoid and squamous phenotypes (**I**). **(J)** Typical epithelial gland-like metastasis in lymph node from *Pten^fl/fl^ Kras^G12V^* cohort. **(K, L)** Rare sarcomatoid nodules in lymph nodes **(K)** combined with sarcomatoid metastases in the lung **(L)** observed in a single mouse (out of 20) in the *Pten^fl/fl^Kras^G12V^* cohort. **(M, N)** Rare small lymph node deposits in *Pten^fl/fl^ Kras^G12V^PlxnB1^-/-^* mouse. **(O)** The single lymph node deposit observed in the *Pten^fl/fl^ Kras^G12V^PlxnB1^WT^* cohort. **(P-R)** Heterogeneous lumbar lymph node metastases from *Pten^fl/fl^ Kras^G12V^PLXNB1^MUT^* mice, including mixed epithelial/sarcomatoid deposits **(P)**, sarcomatoid (**Q**) and squamous metaplasia **(R)**. **(S-V)** Organ metastasis and local invasion in *Pten^fl/fl^ Kras^G12V^PLXNB1^MUT^* mice showing lung metastatic deposit with sarcomatoid (**S**) and squamous histology (**T**), abdominal metastasis adjoining pancreas (**U**) and prostate tumour invading urethra (**V**). Scale bars indicated in individual panels.

Prostate tumours in *Pten^fl/fl^p53^fl/fl^, Pten^fl/fl^p53^fl/fl^PlxnB1^-/-^* and *Pten^fl/fl^p53^fl/fl^PLXNB1^MUT^* models showed similar progression from adenocarcinoma at day 100 towards sarcomatoid metaplasia at 6 months **(Figure 3 A-L)**, metastatic deposits in *Pten^fl/fl^p53^fl/fl^* mice were all represented by the primary sarcomatoid tumours encroaching the lumbar lymph nodes and invading further into the peritoneum (**Figure 3 M-O**, **Supplementary Figure 5)** whereas *Pten^fl/fl^p53^fl/fl^PLXNB1^MUT^* mice had peritoneal and muscle invasion and lymph node metastasis (**Figure 3 P-T, Supplementary Figure 6)**.

**Figure 3.**
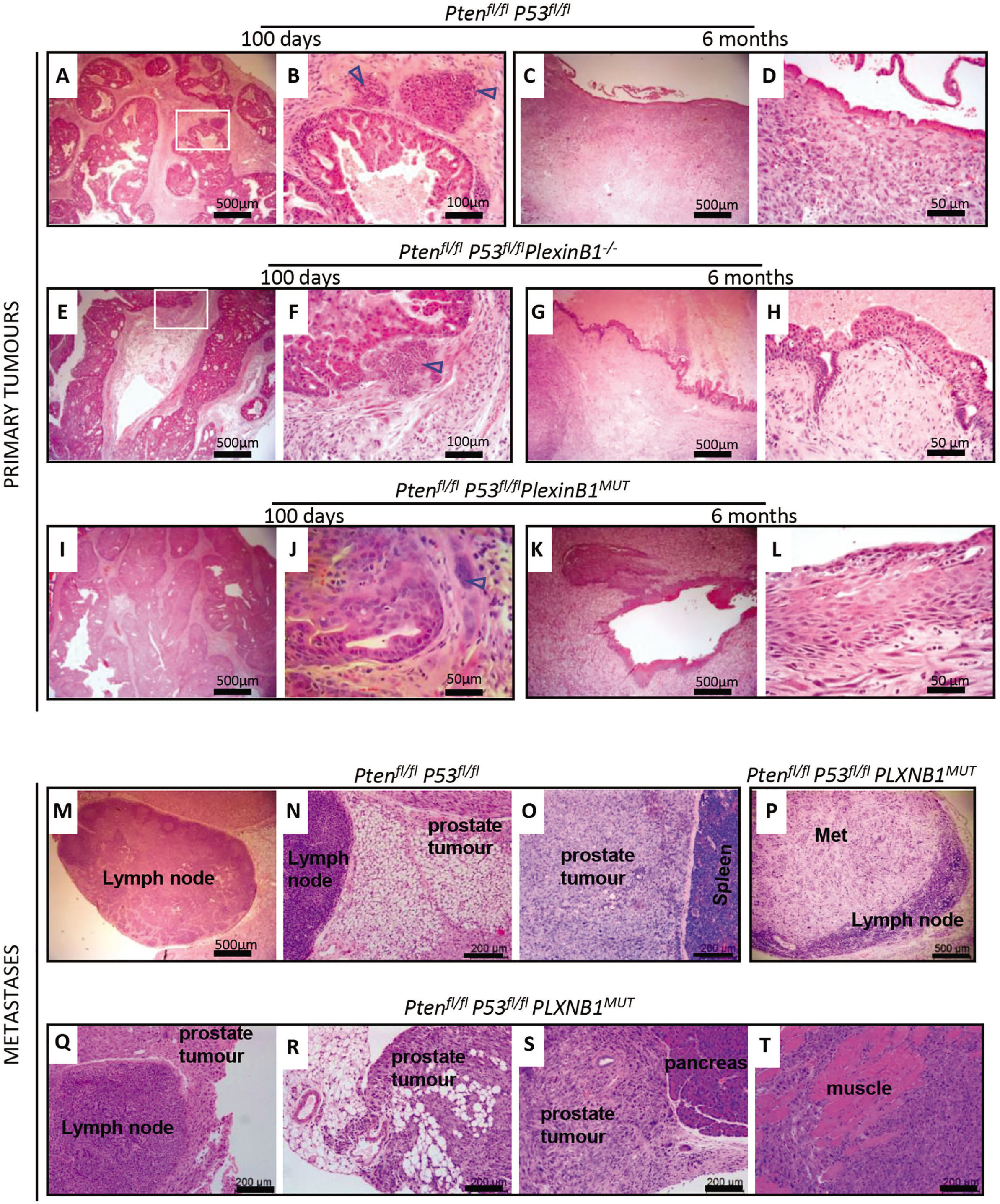
H&E histology of primary prostate tumours and metastatic deposits in *Pten^fl/fl^p53^fl/fl^* cohorts (see also Supplementary Figures 7,8). **(A,B)** Invasive adenocarcinoma in *Pten^fl/fl^ p53^fl/fl^* mouse prostate at day 100 timepoint showing sarcomatoid deposits next to the epithelium (marked with arrows). **(C,D)** Sarcomatoid carcinoma in prostates of 6 month old *Pten^fl/fl^ p53^fl/fl^* mice. **(E,F)** Invasive adenocarcinoma in prostates of *Pten^fl/fl^ p53^fl/fl^ PlxnB1^-/-^* mice at day 100 timepoint showing sarcomatoid deposits next to epithelium (marked with arrow). **(G,H)** Sarcomatoid tumours from prostates of 6 month old *Pten^fl/fl^ p53^fl/fl^ PlxnB1^-/-^* mice. **(I,J)** Invasive adenocarcinoma in mouse prostates at day 100 timepoint in *Pten^fl/fl^ PLXNB1^MUT^* mice; sarcomatoid cells marked by arrows. **(K,L)** Widespread epithelial invasion into stroma and pronounced expansion of sarcomatoid mass combined with squamous differentiation of epithelium in prostates of 7 month old *Pten^fl/fl^ PLXNB1^MUT^* mice. **(M-O)** Prostate sarcomatoid deposits on the perimeter of a lumbar lymph node **(M,N)** and adjoining spleen **(O)** in *Pten^fl/fl^ p53^fl/fl^* mouse. **(P-T)** Metastatic deposits in lumbar lymph nodes **(P)** and on the node perimeter invading to peritoneum **(Q)**, peritoneum **(R)**, pancreas **(S)** and sarcomatoid prostate tumour invading pelvic muscle **(T)** of *Pten^fl/fl^ p53^fl/fl^ PLXNB1^MUT^* mice. Scale bars indicated in individual panels.

*PlxnB1* ablation made no significant difference to survival of either *Pten^fl/fl^Kras^G12V^* or *Pten^fl/fl^p53^fl/fl^* mice; *PLXNB1^WT^* overexpression made no significant difference to survival of the *Pten^fl/fl^Kras^G12V^* line (**Figure 4A,B**). Unexpectedly, however, expression of *PLXNB1^MUT^* significantly increased the survival of both *Pten^fl/fl^Kras^G12V^* and *Pten^fl/fl^p53^fl/fl^* mice (**Figure 4A, B**). *Pten^fl/fl^Kras^G12V^PLXNB1^P1597L^* mice survived for a median of 226.5 days, compared to 182, 187.5 and 176 days for *Pten^fl/fl^Kras^G12V^, Pten^fl/fl^Kras^G12V^PlxnB1^-/-^* and *Pten^fl/fl^Kras^G12V^PLXNB1^WT^* mice respectively (p=0.0148 vs *Pten^fl/fl^Kras^G12V^*, Figure 2A). Prostate tumour growth in *Pten^fl/fl^Kras^G12V^PLXNB1^P1597L^* animals was more heterogeneous. Despite the increased lifespan in this cohort, the animals which had to euthanized by day 200 due to illness had a trend towards earlier development of larger tumours: 7/8 *Pten^fl/fl^Kras^G12V^* mice expressing *PLXNB1^P1597L^* had tumours exceeding 5% of the total mouse weight compared to 4/9 and 5/11 *Pten^fl/fl^Kras^G12V^* and *Pten^fl/fl^Kras^G12V^PlxnB1^-/-^* mice respectively. Median survival of *Pten^fl/fl^p53^fl/fl^PLXNB1^P1597L^* mice was 211 days, compared to 177 days in *Pten^fl/fl^p53^fl/fl^ mice (p<0.001)* and 185 days for *Pten^fl/fl^p53^fl/fl^PlxnB1^-/-^* mice (**Figure 4B**) (p<0.001).

**Figure 4.**
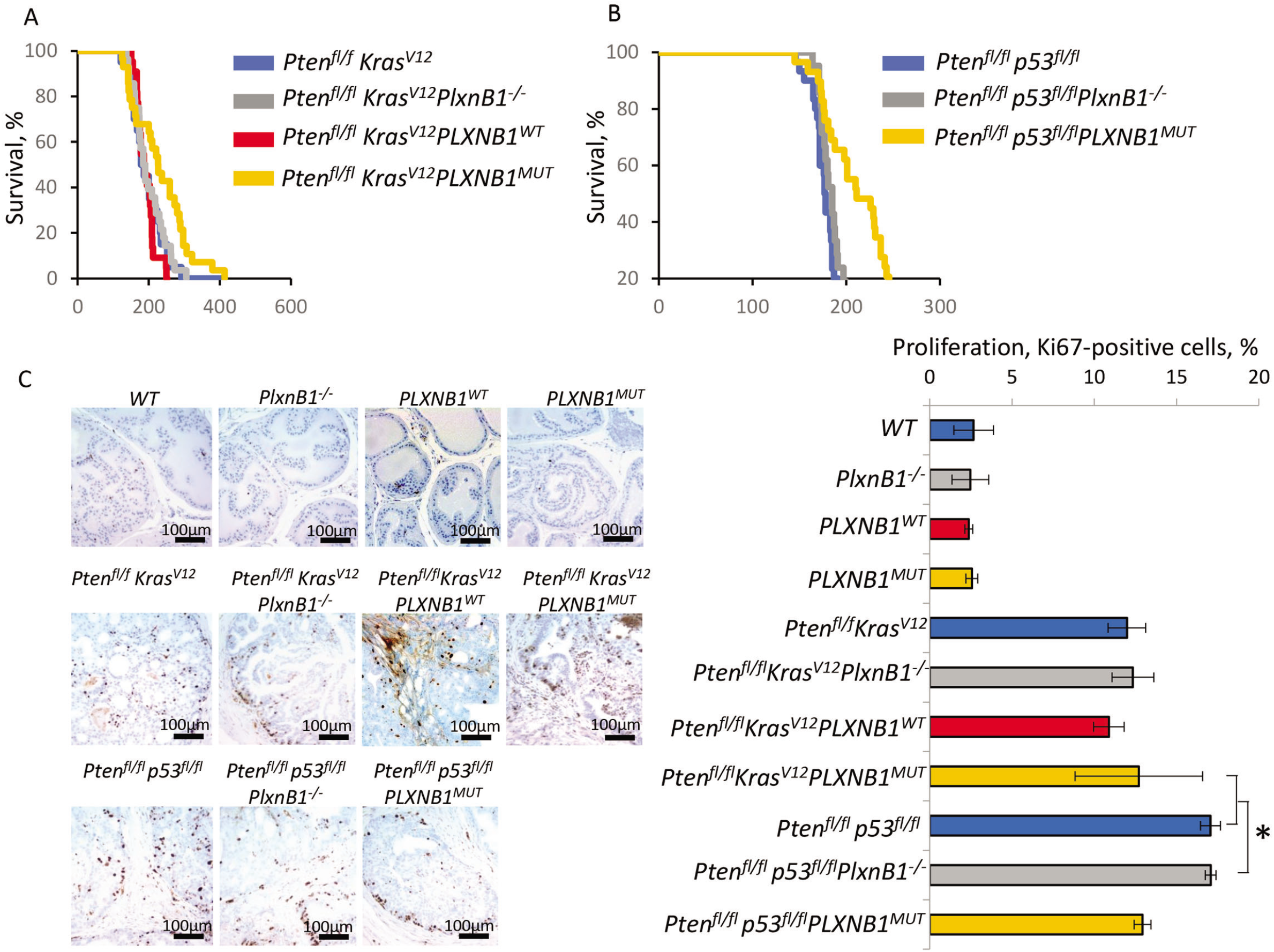
PLXNB1^MUT^ expression suppresses prostate tumour proliferation and extends survival. **(A)** Kaplan-Meier survival curves for *Pten^fl/fl^Kras^G12V^*(n=20), *Pten^fl/fl^Kras^G12V^PlxnB1^-/-^* (n=28), *Pten^fl/fl^Kras^G12V^PLXNB1^WT^*(n=22) and *Pten^fl/fl^Kras^G12V^PLXNB1^MUT^*(n=28) cohorts. Primary prostate tumour growth was the major reason for euthanasia. The increase in survival of the *Pten^fl/fl^Kras^G12V^PLXNB1^MUT^* cohort (median survival 226.5 days) compared to the *Pten^fl/fl^Kras^G12V^cohort* (median 182 days) is significant (log-rank test: z=2.44, p=0.0148, 95% confidence). **(B)** Kaplan-Meier survival curves for *Pten^fl/fl^p53^fl/fl^(n=30), Pten^fl/fl^p53^fl/fl^ PlxnB1^-/-^* (n=21) and *Pten^fl/fl^p53^fl/fl^ PLXNB1^MUT^(n=29)* cohorts. The increase in survival of the *Pten^fl/fl^p53^fl/fl^ PLXNB1^MUT^* cohort (median 211 days) compared to *Pten^fl/fl^p53^fl/fl^* (median 177 days) is significant (log-rank test, z=4.86, p<0.001, 95% confidence). **(C)** *Ki67* antigen staining and quantitation of proliferation rates for prostate epithelium of 100 day old mice from *WT*, *PlxnB1^-/-^ PLXNB1^WT^, PLXNB1^MUT^, Pten^fl/fl^Kras^G12V^, Pten^fl/fl^Kras^G12V^ PlxnB1^-/-^, Pten^fl/fl^Kras^G12V^PLXNB1^WT^, Pten^fl/fl^Kras^G12V^PLXNB1^MUT^, Pten^fl/fl^ p53^fl/fl^, Pten^fl/fl^p53^fl/fl^PlxnB1^-/-^* and *Pten^fl/fl^p53^fl/fl^ PLXNB1^MUT^* mice. *PLXNB1^MUT^* expression suppressed proliferation in the *Pten^fl/fl^p53^fl/fl^* background. *P<0.05 (t-test, n=3, mean±SD). Scale bars = 100μm.

Consistent with these findings, deletion of PlexinB1 had a negligible effect on cell proliferation in either model at 100 days, as demonstrated by Ki67 staining of prostate epithelial cells. Cell proliferation rates in *Pten^fl/fl^Kras^G12V^PLXNB1^WT^* tumours were also not statistically different from those of *Pten^fl/fl^Kras^G12V^* cohorts (**Figure 4C**). However, there was a wide variation in tumour cell Ki67 staining between different *Pten^fl/fl^Kras^G12V^PLXNB1^MUT^* mice at 100 days, suggesting that this cohort developed a heterogeneous mix of slowly and rapidly growing primary tumours which overall resulted in an increase in median survival. The results for the *Pten^fl/fl^p53^fl/fl^* background mice were more consistent, with Ki67 staining of prostates of 100 day *Pten^fl/fl^p53^fl/fl^PLXNB1^MUT^* mice showing a 1.32-fold decrease in cell proliferation compared to the *Pten^fl/fl^p53^fl/fl^ cohort* (p<0.01) (**Figure 4C**). As prostate cancer mouse models typically need to be euthanized as a result of local complications associated with primary tumour bulk, the suppression of primary tumour proliferation by *PLXNB1^MUT^* overexpression explains the extended survival in these models.

### PlexinB1^MUT^ increases metastasis in mouse models of prostate cancer

Next, we quantified the metastatic lesions in the different tumour cohorts by histological examination and staining for androgen receptor (AR), a prostate epithelial cell marker, to confirm the prostate origin of metastatic lesions (**Figure 2J-O; Figure 3M-T; Supplementary Figures 2-6**). Genetic deletion of PlexinB1 reduced metastasis substantially in both *Pten^fl/fl^Kras^G12V^* and *Pten^fl/fl^p53f^fl/fl^* models (p=0.0411 for *Pten^fl/fl^Kras^G12V^mice*, **Table 1, Figure 5**). Deletion of Plexin-B1 resulted in a three-fold reduction in the number of mice with metastases in the *Pten^fl/fl^Kras^G12V^* model: 35% of *Pten^fl/fl^Kras^G12V^* mice had node metastases, including one mouse with an additional lung metastasis; in contrast, 10.7% of *Pten^fl/fl^Kras^G12V^PlxnB1^-/-^* mice had node metastases and no distant metastases were found (**Figure 5A,B**). Deletion of *PlxnB1* in *Pten^fl/fl^p53^fl/fl^* mice completely blocked metastases (**Figure 5C,D**). Remarkably, overexpression of *PLXNB1^WT^* also significantly suppressed the metastatic spread of tumours in *Pten^fl/fl^Kras^G12V^mice* (p=0.0121 χ^2^ test), with only a single local metastasis observed in *Pten^fl/fl^Kras^G12V^PLXNB1^WT^mice* (4.5%) (**Figure 5A,B**).

**Figure 5.**
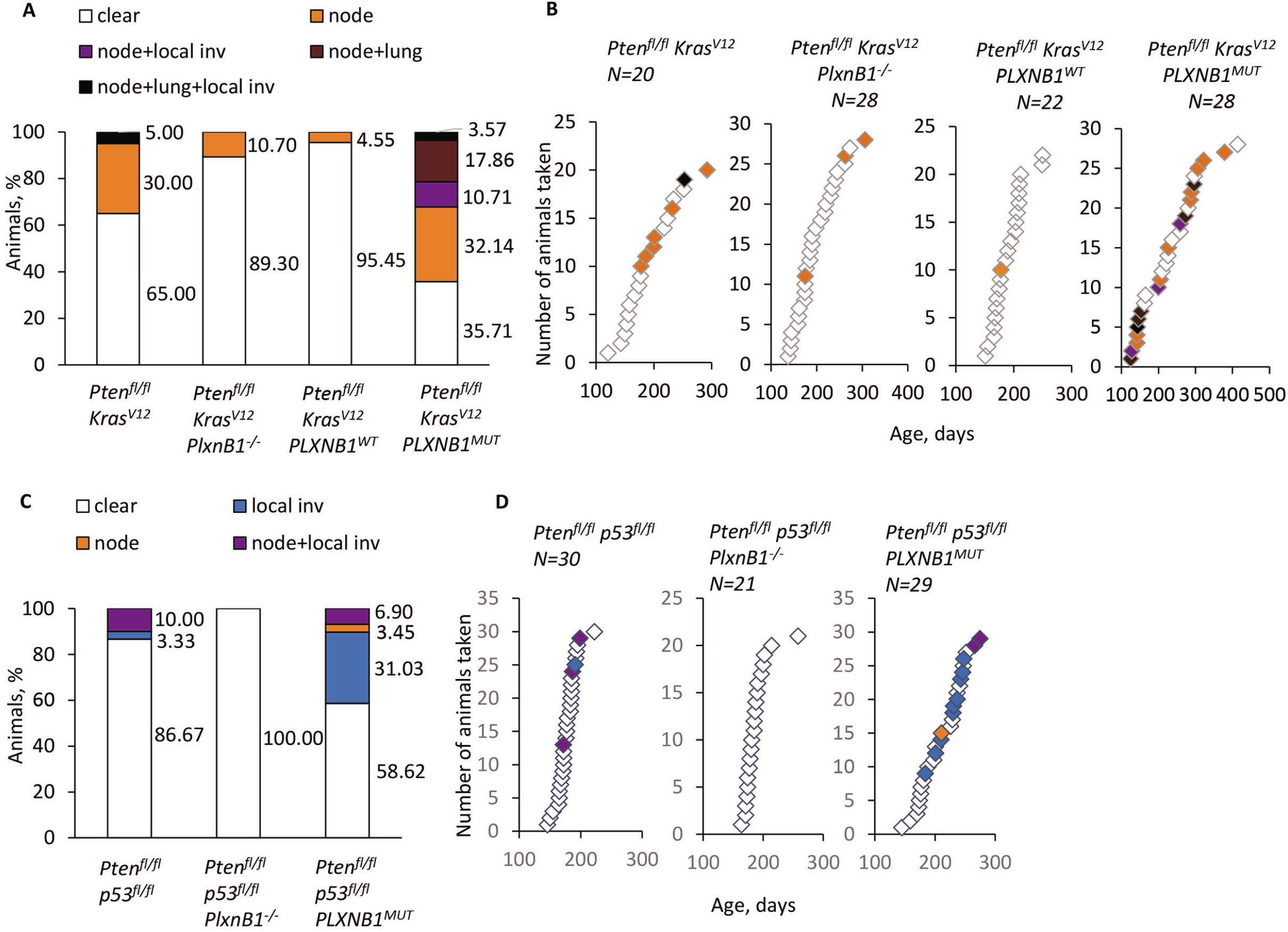
PLXNB1^MUT^ increases metastasis whereas PlexinB1 deletion or PLXNB1^WT^ expression suppress metastasis in mouse models of prostate cancer. Following necropsy, mice were categorised according to their metastatic outcome: no metastatic deposits (white), lymph node metastasis (orange), invasion into peritoneum or pelvic muscle (blue), lymph node metastasis combined with invasion into peritoneum or pelvic muscle (purple), combined lymph node and lung metastasis (brown), animals with both lymph node and lung metastasis combined with invasion into peritoneum or pelvic muscle (black). **(A)** Percentages of animals affected/not affected by metastasis in *Pten^fl/fl^Kras^G12V^* cohorts. **(B)** Timing and type of metastatic deposits in *Pten^fl/fl^Kras^G12V^* cohorts. **(C)** Percentage of animals affected/not affected by metastasis in *Pten^fl/fl^ p53^fl/fl^* cohorts. **(D)** Timing and type of metastatic deposits in *Pten^fl/fl^ p53^fl/fl^* cohorts. See **Table 1** for statistical analyses.

**Table 1.**
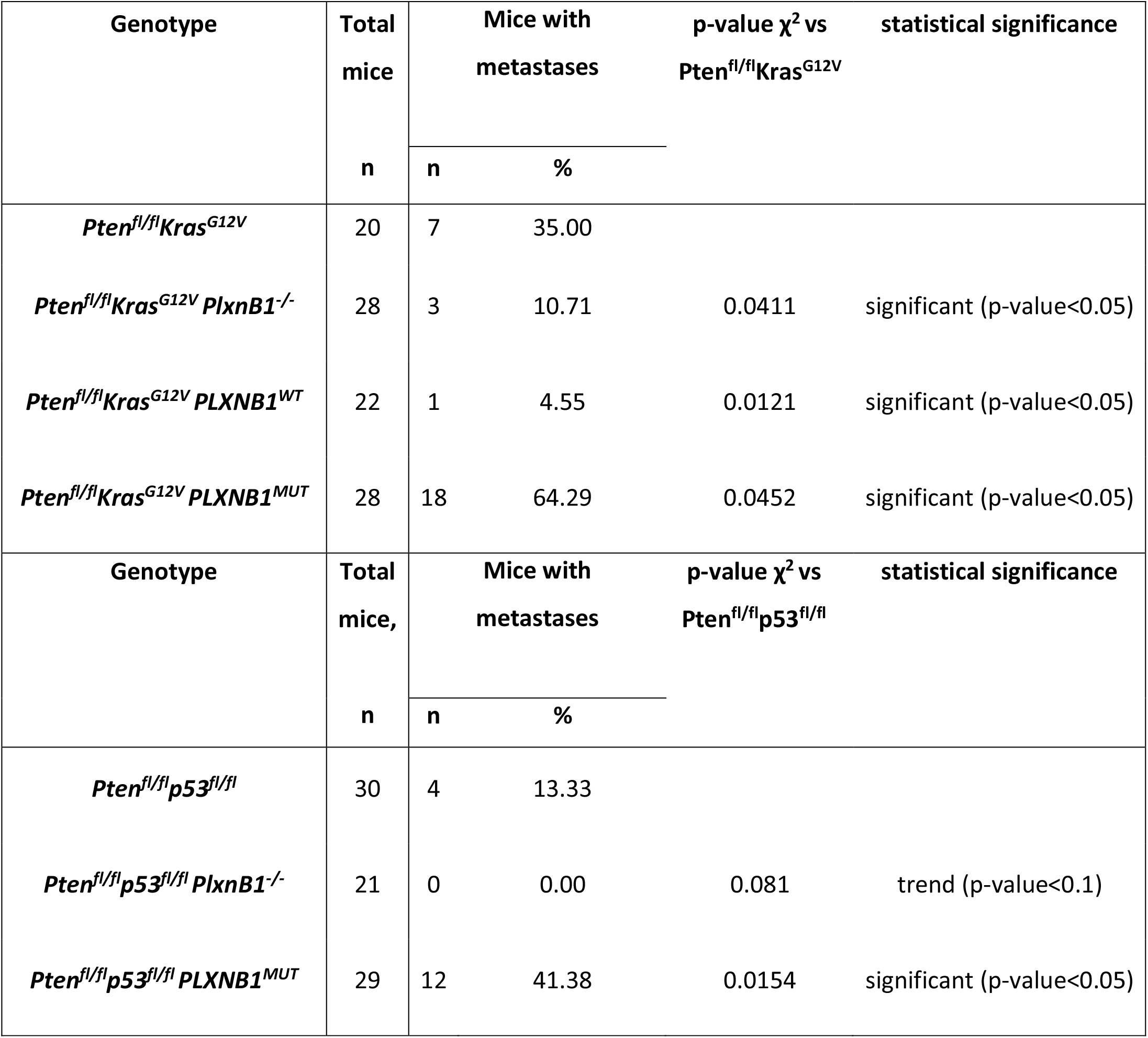
Number of mice in each cohort, numbers of mice with metastases and statistical significance of differences compared to control.

Importantly, however, expression of mutant *PLXNB1^MUT^* in both *Pten^fl/fl^Kras^G12V^* and mice *Pten^fl/fl^p53^fl/fl^* significantly increased the percentage of mice with metastases, compared to the parental lines (p=0.0452 vs *Pten^fl/fl^Kras^G12V^* and p=0.0154 vs *Pten^fl/fl^p53^fl/fl^* respectively), to *Pten^fl/fl^Kras^G12V^PlxnB1^-/-^* and *Pten^fl/fl^p53^fl/fl^PlxnB1^-/-^* mice (p<0.001 for both lines) and to *Pten^fl/fl^Kras^G12V^PLXNB1^WT^* mice (p<0.0010; **Figure 5, Table 1**). On the *Pten^fl/fl^Kras^G12V^* background, this increase in metastases was particularly evident in sites distant from the prostate - the number of mice with lung metastases increased from 5% to 21.43% upon expression of *PLXNB1^MUT^*, whereas no lung metastases were found for *Pten^fl/fl^Kras^G12V^PlxnB1^-/-^* or *Pten^fl/fl^Kras^G12V^PLXNB1^WT^* mice. On the *Pten^fl/fl^p53^fl/fl^* background, *PLXNB1^MUT^* mice showed a marked increase in locally invasive tumours demonstrating invasion of the primary tumour into local structures or organs including peritoneum, pelvic or bladder muscle, vas deferens, and lymph node (**Supplementary Figure 6**).

### PlexinB1 increases local invasion of prostate tumour cells

Our results show that ablation of the *PlxnB1* gene and prostate epithelial cell-specific overexpression of wild-type Plexin-B1 both decrease metastasis, while overexpression of mutant Plexin-B1 has the opposite effect. Cancer metastasis is a multistep process which begins with tumour cell invasion from prostate acini through the basal membrane and into the prostate stroma. To establish the stage at which wild-type or mutant Plexin-B1 expression affects metastasis, we investigated the effect of expression of the different forms of Plexin-B1 on invasion of primary tumour cells into the prostate stroma.

Primary tumours from all cohorts of *Pten^fl/fl^Kras^G12V^ or Pten^fl/fl^p53^fl/fl^* mice were immunostained for the epithelial marker pan-cytokeratin at the early timepoint of 100 days (before the onset of metastasis) and the percentage of tumours breaking into the stroma was scored (**Figure 6**). Deletion of Plexin-B1 significantly reduced invasion in both *Pten^fl/fl^Kras^G12V^* (2.6% to 0.7%, p>0.05) and *Pten^fl/fl^p53^fl/fl^* (1.6% to 0.4% p>0.05) backgrounds, suggesting that Plexin-B1 enhanced invasion in both models. Overexpression of wild-type Plexin-B1 in prostate epithelial cells of *Pten^fl/fl^Kras^G12V^* mice also suppressed tumour cell invasion into the stroma (p>0.05). In contrast, overexpression of mutant Plexin-B1 in prostate epithelial cells increased invasion in both *Pten^fl/fl^Kras^G12V^* and *Pten^fl/fl^p53^fl/fl^* models (2.6% to 8% p>0.05, and 1.6% to 2.3% p>0.05 respectively). These results show that mutant Plexin-B1 enhances metastasis and wild-type PlexinB1 inhibits metastasis at an early stage in the metastatic process.

**Figure 6.**
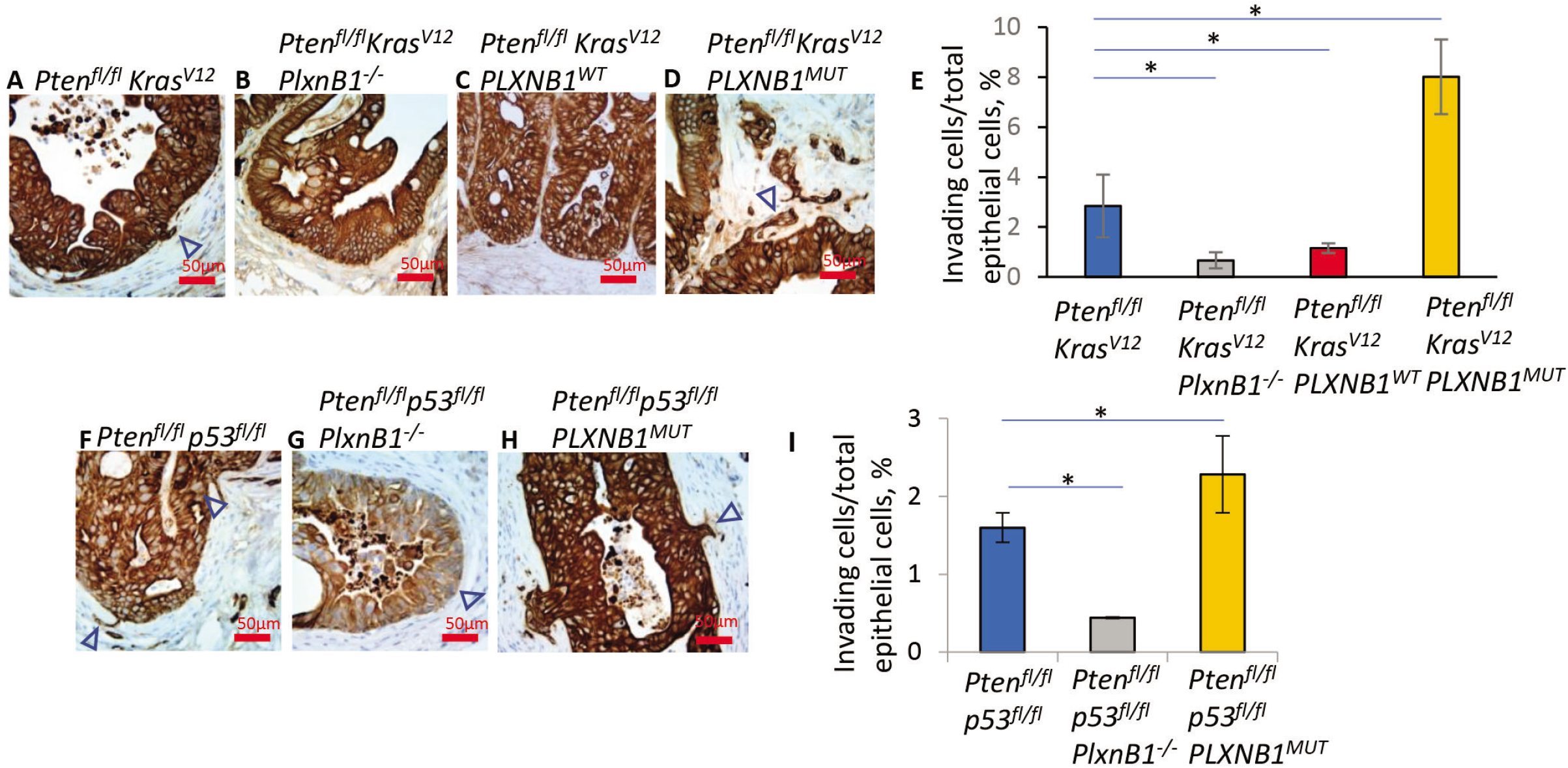
PLXNB1^MUT^ expression promotes local invasion by prostate tumour cells. Immunostaining of prostates of 100 day old mice with cytokeratin AE1/AE3 (pan-cytokeratin) to identify prostate epithelial cells breaking basement membrane and invading stroma *in Pten^fl/fl^Kras^G12V^* **(A-E)** and *Pten^fl/fl^ p53^fl/fl^* cohorts **(F-I)**. Invading cells are indicated with arrowheads. Scale bars are 50μm. Quantitation in the *Pten^fl/fl^Kras^G12V^* and *Pten^fl/fl^ p53^fl/fl^* backgrounds shown in **(E)** and **(I)** respectively. Pan-cytokeratin positive cells breaking the basement membrane or located inside the stromal compartment were counted and divided by total number of pan-cytokeratin positive cells. *P<0.05 (t-test, n=3, mean±SD).

Overall, these findings support a model in which expression of mutant Plexin-B1 switched prostate tumour cells from a proliferative to an invasive phenotype. This led to longer survival of the mice, as there were fewer local complications due to primary tumour mass, but increased tumour cell motility, escape from the primary tumour site and metastasis to lymph nodes and organs. It is important to note that men with prostate cancer rarely, if ever, die of complications of the primary tumour (unlike with mice), but rather of the metastatic burden.

### PLXNB1^MUT^ expression correlates with Rho/ROCK pathway activation in mouse prostate tumours

Plexin-B1 activates RhoA and RhoC through PDZRhoGEF/LARG(12), which bind to the C-terminus of Plexin-B1 and inactivates RhoA and RhoC through p190RhoGAP and Rap inactivation(32, 33). Plexin-B1-mediated activation of RhoA/C is a key pathway promoting metastasis in ErbB2-mouse models of breast cancer(18). To establish if Plexin-B1 might signal via RhoA/C to promote metastasis in the mouse models of prostate cancer, phosphorylation of myosin light chain (MLC2) phospho-MLC2^Ser19^ (a marker of Rho-associated protein kinase (ROCK) activation(34)) was evaluated in tumours of *Pten^fl/fl^Kras^G12V^* and *Pten^fl/fl^p53^fl/fl^* models (**Figure 7**). Deletion of Plexin-B1 reduced semi-quantitative MLC2 phosphorylation H-score, with a two-fold reduction in cells with ‘strong’ staining in both models upon PlexinB1 ablation in 100 day old mice (p= <0.05, **Figure 7H**). Overexpression of wild-type Plexin-B1 in the *Pten^fl/fl^Kras^G12V^* model also decreased MLC2 phosphorylation in tumours (p=<0.05; **Figure 7H**).

**Figure 7.**
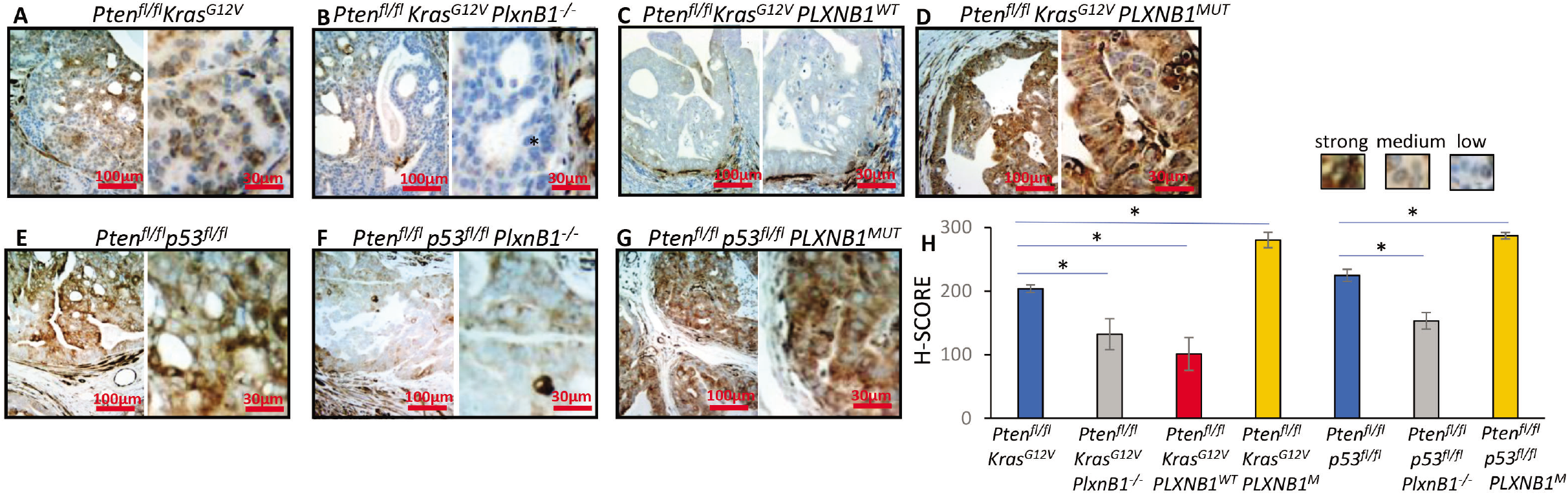
PLXNB1^MUT^ expression in the mouse prostate epithelium stimulates myosin phosphorylation. Immunostaining of mouse prostates for phospho-Myosin Light Chain 2 Ser19 (phospho-MLC2^Ser19^) to identify levels of cell contractility and Rho-kinase (ROCK) activation in *Pten^fl/fl^Kras^G12V^* cohorts at 100 days **(A-D)** and *Pten^fl/fl^p53^fl/fl^* cohorts at 100 days **(E-G)**. Relative to *PlxnB1* intact controls **(A,E)** *PlxnB1* deletion leads to lower staining intensity both in *Pten^fl/fl^Kras^G12V^* **(B)** and *Pten^fl/fl^p53^fl/fl^* cohorts **(F)**. *PLXNB1^WT^* expression lowers MLC2 phosphorylation in the *Pten^fl/fl^Kras^G12V^* cohort **(C)**. *PLXNB1^MUT^* expression increases MLC2 phosphorylation on both backgrounds **(D, G)**. Scale bars are 100 μm (left image) and 30 μm (right image). **(H)** H-score quantitatation of phospho-MLC2^Ser19^ staining. Epithelial cells were divided into three categories according to staining intensity (strong/medium/low). H-score = 1 × (% «low staining» cells) + 2 × (% «medium staining» cells) + 3 × (% «strong staining» cells). Cells were counted in all seven cohorts (n=3, 5 fields per sample, t-test). *P<0.05 (t-test, n=3, mean±SD).

In contrast, expression of *PLXNB1^MUT^* significantly increased MLC2 phosphorylation H-score in both *Pten^fl/fl^Kras^G12V^* and *Pten^fl/fl^p53^fl/fl^* models (4.8 fold and 3.3 fold increase in the percentage of cells with strong staining for *Pten^fl/fl^Kras^G12V^PLXNB1^P1597L^* and *Pten^fl/fl^p53^fl/fl^PLXNB1^P1597L^* respectively, p=<0.05, **Figure 7H**). These results suggest that PlexinB1 promotes metastasis at least in part, via the Rho-ROCK pathway.

## Discussion

Our results show that PlexinB1 status has a major effect on prostate cancer metastasis. Deletion of PlexinB1 in *Pten^fl/fl^Kras^G12V^* and *Pten^fl/fl^p53^fl/fl^* mice significantly decreased metastasis in comparison to *Pten^fl/fl^Kras^G12V^* and *Pten^fl/fl^p53^fl/fl^* mice with normal levels and patterns of PlexinB1 expression, demonstrating that Plexin-B1 is required for metastasis in these mouse models. Plexin-B1 deletion used a germline, whole body knockout approach, however, and Plexin-B1 is normally expressed by endothelial cells in the tumour microenvironment in addition to tumour epithelial cells (for example, see **Figure 1H**). Activation of Plexin-B1 on endothelial cells by Sema4D expressed on prostate tumour cells promotes angiogenesis(35), therefore whole body knockout of Plexin-B1 in our mouse models would inhibit Sema4D-induced angiogenesis and may also contribute to the decrease in metastasis observed. Consistent with these results, whole body knockout of PlexinB1 inhibited metastasis in ErbB2-expressing models of breast cancer(18). In contrast, xenografts of melanoma cells expressing Sema4D and Plexin-B1 in Plexin-B1 knockout mice still develop tumours(36). Furthermore, knockdown of Plexin-B1 in prostate cancer cells expressing ErbB2 reduces motility and invasion *in vitro*, consistent with the *in vivo* results(37) and supporting a tumour cell-specific role for Plexin-B1.

Indeed, overexpression of wild-type Plexin-B1 targeted specifically to the prostate epithelial cells of *Pten^fl/fl^Kras^G12V^* mice, decreased invasion and metastasis in comparison to *Pten^fl/fl^Kras^G12V^* mice expressing normal levels of Plexin-B1. Sema4D is expressed by cells in the tumour stroma and tumour-associated macrophages(35) and Sema4D secreted from the tumour microenvironment may act as a repellent cue to inhibit migration and invasion of tumour cells expressing wild-type Plexin-B1, confining the tumour cells to the primary tumour mass. Sema4D produced by tumour cells may also act as an autocrine or paracrine signal to suppress migration; non polarised activation of Plexin-B1 over the whole cell results in cell collapse *in vitro(38)*. Alternatively, ligand independent Plexin-B1 signalling due to wild-type Plexin-B1 overexpression and receptor clustering(39), may repress migration and invasion.

Interestingly, while high levels of wild-type Plexin-B1 in prostate epithelial cells decreased invasion and metastasis in *Pten^fl/fl^Kras^G12V^* mice, similar levels of mutant (P1597L) Plexin-B1 significantly increased metastasis. These findings reflect our previous results where overexpression of wild-type Plexin-B1 decreased migration and invasive capacity of HEK293 cells and overexpression of the P1597L mutant form of Plexin-B1 increased motility and invasion(19). The increase in invasion and metastasis of prostate cancers observed in the *Pten^fl/fl^Kras^G12V^PLXNB1^MUT^* (**Figures 5,6**) mice may result from a change in response of tumour cells to semaphorins produced by the stroma – a switch from repulsion to attraction(4). Sema4D has been shown to promote or suppress migration and invasion depending on cellular context and the Plexin-B1 co-receptors expressed by the responding cell(5).

The contrasting results from overexpression of the wild type and mutant proteins argues against the findings being an artefact of overexpression *per se*. Rather, the complexity of the outcomes of the different cohorts likely results from the multiple mechanisms through which Plexin-B1 can activate its downstream effectors. Wild type Plexin-B1 regulates several small GTPases including Rap(16) which, like most GTPases, cycles between an active GTP-bound and an inactive GDP-bound form. Rap has diverse functions in tumour progression(40) and Rap1 activation promotes prostate cancer metastasis(41). GTP-bound Rap activates Ras(42), Rac(43) and Rho(44) signalling enhancing tumour growth, cell motility and invasion. Wild-type PlexinB1 acts as a GTPase activating protein (GAP) for Rap(16), catalysing the conversion of RapGTP to inactive RapGDP. WT PlexinB1, through its activity as a RapGAP, thereby reduces Rap activation, decreasing Ras, Rac, and Rho activity and suppressing cell motility, invasion and MLC2 phosphorylation (**Supplementary Figure 7**). The P1597L mutation occurs in the GAP domain of PlexinB1 and is predicted to disrupt the GAP activity of PlexinB1. Consequently, RapGTP is not inactivated by mutant PlexinB1, allowing Rap in its active GTP bound form to promote Ras, Rho and Rac activation leading to MLC2 phosphorylation, invasion and metastasis. In this way, the P1597L mutation would result in a loss of inhibitory function (**Supplementary Figure 7**). However, both the WT and mutant form of PlexinB1 also activate Rho through PDZRhoGEF/LARG(12) binding to the C-terminal of PlexinB1; this is not affected by the P1597L mutation. Therefore, overexpression of mutant PlexinB1 results in strong PDZRhoGEF-dependent Rho activation while no longer inhibiting RapGTP, overexpression of wild-type Plexin-B1 results in strong PDZRhoGEF-dependent Rho activation but with inhibition of the RapGTP-dependent Ras/Rho/Rac pathway and Plexin-B1 deletion prevents PDZRhoGEF-dependent Rho activation whilst allowing RapGTP-dependent Ras/Rho/Rac activation. As only mutant PlexinB1 promoted metastasis, and the other two conditions inhibited it, this implies both routes of Rho activation are necessary for maximal metastatic potential. However, we cannot exclude the possibility that the mutation may also confer an additional gain of function by an unknown mechanism. We are currently generating new cohorts of Plexin-B1 knockout mice with conditional deletion of PDZRhoGEF or RhoA/C to further understand these mechanisms.

Expression of mutant Plexin-B1 in *Pten^fl/fl^Kras^G12V^* mice resulted in tumours with an epithelial phenotype and a significant increase in metastasis to lymph nodes and lung, metastasis in this background occurring via the lymphatic and/or via blood vessel route. In contrast, expression of mutant Plexin-B1 in *Pten^fl/fl^p53^fl/fl^* mice resulted in an increase in tumours with a mesenchymal phenotype which were predominantly locally invasive into surrounding tissues such as the peritoneum, pelvic or bladder muscle or vas deferens. Consistent with these results, Sema3C drives epithelial mesenchymal transition (EMT) in prostate cells(45) promoting a spindle-like morphology. These results demonstrate the context dependence of specific semaphorin/plexin mediated signalling pathways controlling metastasis.

Plexin-B1 is overexpressed in some cancers and appears to act as a tumour suppressor gene in others(46). Indeed, high levels of Plexin-B1 expression predict longer overall survival in bladder carcinoma, head and neck squamous cell carcinoma and kidney papillary renal cell carcinoma but shorter overall survival in thymoma and kidney renal clear cell carcinoma (data from K-M Plotter Pan-Cancer Tool; https://kmplot.com/analysis/index.php?p=background)(47). Data on the prognostic significance of Plexin-B1 in prostate cancer are complicated by the use of different baseline comparators. A large scale gene expression comparison between prostate cancer and normal tissue(27) showed Plexin-B1 expression was altered in 30% of prostate cancer patients (z-score=±2) and Plexin-B1 expression downregulation was 3 times more common (22.67%) in prostate cancer than its increase (7.33%). However, other large scale genomics projects using a different baseline (diploid tumour samples instead of normal prostate tissue) have suggested that Plexin-B1 upregulation is more common than its decrease. These studies are available through cBioportal(48, 49) and summarised in **Supplementary Table 1**.

Furthermore, the genomic data from cBioportal (http://cbioportal.org)(48, 49) suggests that Plexin-B1 mutation is more detrimental to survival than amplification or deletion; patients with Plexin-B1 mutation had a reduced lifespan (median survival=37 months, 14 cases, 4 deceased) compared to patients with Plexin-B1 copy number alterations (median survival=109 months, 16 cases, 6 deceased, log-rank p-value<0.05) and to the unaltered group (median survival=156 months, 1445 cases, 142 deceased, log-rank p-value<0.05). Notably, cBioportal data also shows that Plexin-B1 expression is inversely correlated with PTEN (Spearman’s correlation −0.374, p-value=1.34E-05, q-value=2.56E-05). It interesting to speculate that elevated levels of wild-type Plexin-B1 may suppress metastasis in a *PTEN*-deleted tumour, but if the Plexin-B1 acquires a mutation it may switch to a driver of aggressive metastatic disease.

Human prostate tumours are highly heterogenous and consist of a complex mixture of clones of different genetic make-up which complicates analysis(50). The use of mouse models of a defined genetic background allows the effect of the many different genetic changes found in human tumours to be analysed separately. Current treatments for metastatic prostate cancer are effective only in the short term, highlighting the need for new therapies for late stage disease. In order to test such therapies, pre-clinical models in which metastasis is driven by clinically relevant mutations, such as those we have developed here, are a key requirement. Indeed, our results have demonstrated that PlexinB1 has a complex yet significant role in metastasis and is a potential therapeutic target to block the lethal spread of prostate cancer.

## Author Contributions

Study conception and design: BS, MW, NT, TW, SO, JM, ARC. Acquisition of data: BS, NT. Analysis and interpretation of data: BS, MW, NT, DG, MJS, ARC. Drafting of manuscript: BS, MW, MJS.

## Acknowledgements

This work was funded by Prostate Cancer Research Centre. We also thank Elaine Taylor for assistance with mouse husbandry and Derek Scarborough and Marc Isaacs for their assistance in histology. This manuscript is dedicated to the memory of the late Professor Alan Clarke. This manuscript has been deposited as a preprint at bioRxive (https://biorxiv.org/cgi/content/short/2020.07.19.203695v1).

