## Supplementary Figure 1 for "Plexin-B1 mutation drives prostate cancer metastasis"

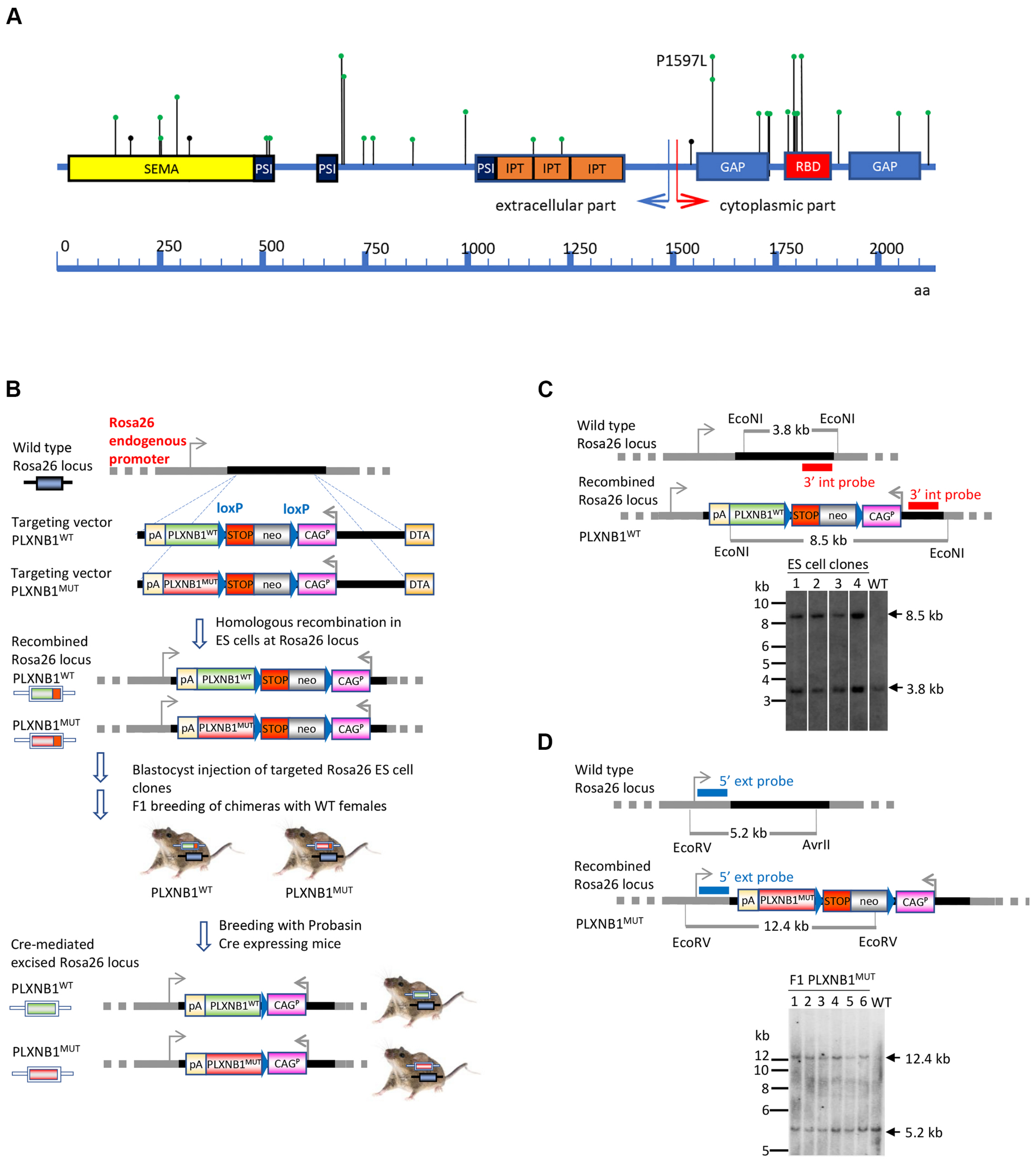
A
P1597L
SEMA
PSI
PSI
PSI
IPT
IPT
IPT
GAP
RBD
GAP
extracellular part
cytoplasmic part
0
250
500
750
1000
1250
1500
1750
2000
aa
B
Wild type Rosa26 locus
Rosa26 endogenous promoter
loxP
loxP
Targeting vector PLXNB1WT
pA
PLXNB1WT
STOP
neo
CAGP
DTA
Targeting vector PLXNB1MUT
pA
PLXNB1MUT
STOP
neo
CAGP
DTA
Homologous recombination in ES cells at Rosa26 locus
Recombined Rosa26 locus
PLXNB1WT
PLXNB1MUT
Blastocyst injection of targeted Rosa26 ES cell clones
F1 breeding of chimeras with WT females
PLXNB1WT
PLXNB1MUT
Breeding with Probasin Cre expressing mice
Cre-mediated excised Rosa26 locus
PLXNB1WT
PLXNB1MUT
C
Wild type Rosa26 locus
EcoNI
3.8 kb
EcoNI
3' int probe
Recombined Rosa26 locus
3' int probe
PLXNB1WT
8.5 kb
EcoNI
EcoNI
ES cell clones
kb
1 2 3 4 WT
10
8
6
5
4
3
8.5 kb
3.8 kb
D
Wild type Rosa26 locus
5' ext probe
5.2 kb
EcoRV
AvrII
Recombined Rosa26 locus
5' ext probe
PLXNB1MUT
12.4 kb
EcoRV
EcoRV
F1 PLXNB1MUT
1 2 3 4 5 6 WT
kb
12
10
8
6
5
12.4 kb
5.2 kb

**Supplementary Figure 1**. **Generation of *PLXNB1*$^{WT}$ and *PLXNB1*$^{MUT}$ mice. (A)** Schematic representation of the human PLXNB1 protein with its functional domains and distribution of the known cancer-associated mutations. Protein domains: SEMA, sema domain, PSI, domain found in plexins, semaphorins, integrins; IPT, immunoglobulin-like fold shared by plexins and transcription factors; GAP, GTPase activating protein domain, RBD, Rho GTPase binding domain. PLXNB1 mutations are shown by green (missense) or black (frame shift or splice mutations) dots with arms. Higher incidence mutations are indicated by taller arms. P1597L mutation used in this study is located in the GAP domain. **(B)** Schematic representation of *PLXNB1*$^{WT}$ and *PLXNB1*$^{MUT}$ mouse generation based on *Rosa26* targeting strategy. The models were generated by targeted insertion of the *PLXNB1*$^{WT}$ or *PLXNB1*$^{MUT}$ cDNA within the *Rosa26* locus via homologous recombination in embryonic stem cells to establish *PLXNB1*$^{WT}$ or *PLXNB1*$^{MUT}$ cDNA expression under the control of the ubiquitous exogenous *CAG* promoter. A *loxP*-flanked (floxed) transcriptional *STOP* cassette was incorporated between *PLXNB1*$^{WT}$ or *PLXNB1*$^{MUT}$ cDNA and *CAG* promoter to allow transgene expression only in the presence of Probasin (*Pb) Cre* recombinase. Scheme is not depicted to scale. Dotted and solid line represents intronic sequence located in the endogenous *Rosa26* locus and in the targeting vector respectively. *DTA* negative selection cassette is marked in yellow, the *CAG* promoter is shown in pink, *loxP* sites are presented as blue triangles, the combined *STOP-neomycin* selection cassette is indicated by a grey box, the mutant *PLXNB1*$^{WT}$ cDNA is shown in green and the *PLXNB1*$^{MUT}$ cDNA is in red and *hGH polyA* is indicated in beige. **(C)** Southern blot confirmation of presence of *PLXNB1*$^{WT}$ cassette in ES cell clones (4 heterozygous clones identified by PCR). The genomic DNA of ES clones was compared to wild type DNA (C57BL/6). Genomic DNA samples were digested with EcoNI, blotted on nylon membrane and hybridised with an internal 3` probe (indicated in red). This detected a 3.8 kb fragment in wild type genomic DNA and a 8.5 kb hybridisation signal in recombined DNA. None of the positive ES cell clones contain an additional randomly integrated copy of the targeting construct. **(D)** Southern blot confirmation of presence of *PLXNB1*$^{MUT}$ cassette in F1 generation (6 heterozygous mice identified by PCR). The genomic DNA of the F1 mice was compared to wild type DNA (C57BL/6). Genomic DNA samples were digested with AvrII/EcoRV, blotted on nylon membrane and hybridised with an external 5` probe. This detected a 5.2 kb fragment in wild type genomic DNA and a 12.4 kb hybridisation signal in recombined DNA.
