## Supplementary Figure 2 for "Plexin-B1 mutation drives prostate cancer metastasis"

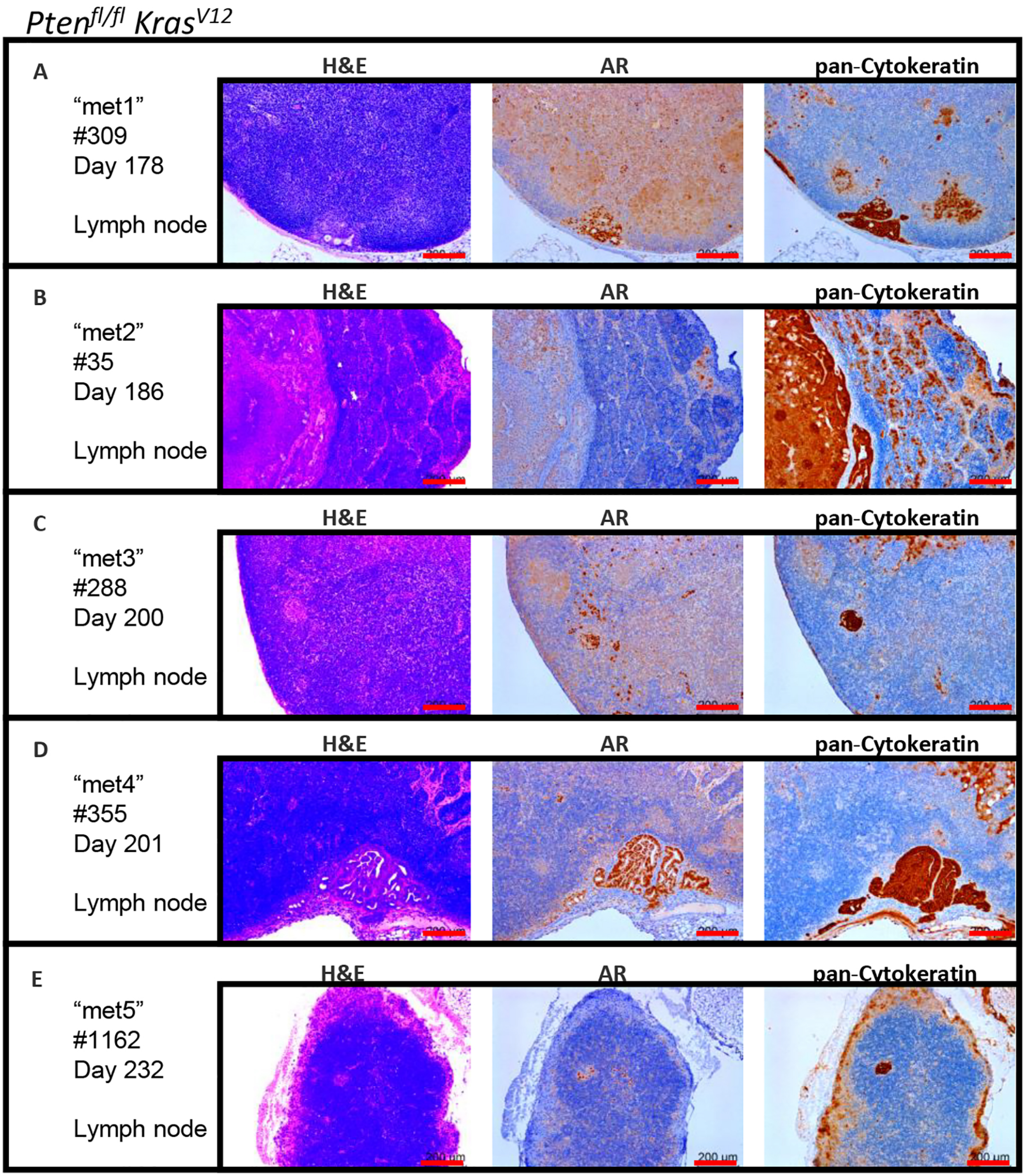

$Pten^{fl/fl}$ $Kras^{V12}$
A
"met1"
#309
Day 178
Lymph node
B
"met2"
#35
Day 186
Lymph node
C
"met3"
#288
Day 200
Lymph node
D
"met4"
#355
Day 201
Lymph node
E
"met5"
#1162
Day 232
Lymph node
H&E
AR
pan-Cytokeratin
200 um

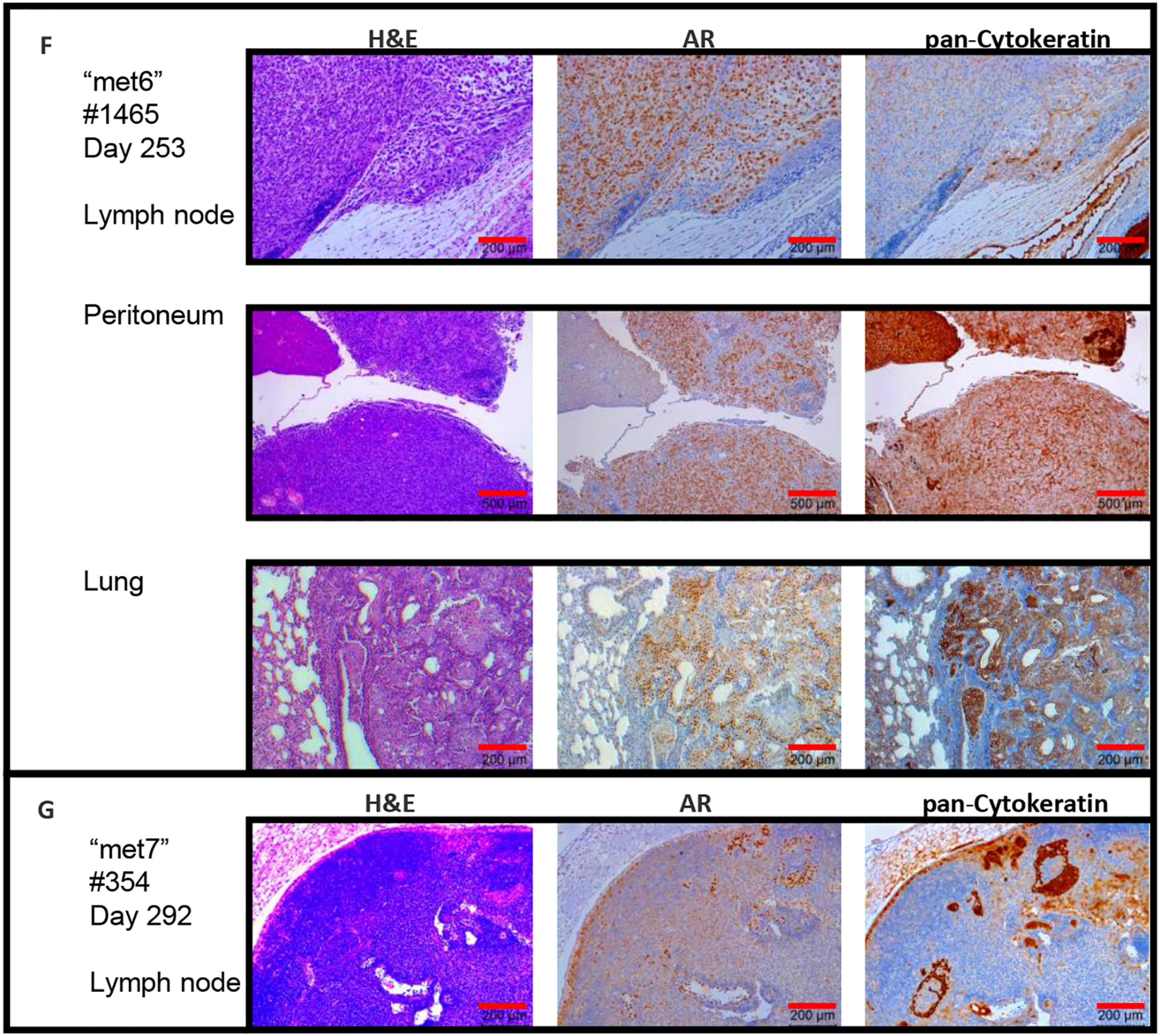

F
H&E
AR
pan-Cytokeratin
"met6"
#1465
Day 253
Lymph node
Peritoneum
Lung
200 µm
500 µm
G
H&E
AR
pan-Cytokeratin
"met7"
#354
Day 292
Lymph node
200 µm

**Supplementary Figure 2. Metastatic deposits in *Pten*$^{fl/fl}$*Kras*$^{G12V}$ mouse cohort stained for H&E, androgen receptor (AR) and pan-cytokeratin.** Metastatic deposits were observed in 7 *Pten*$^{fl/fl}$ *Kras*$^{V12}$ cohort animals, met1 (#309, 178 days old, **A**), met2 (#35, 186 days old, **B**), met3 (#288, 200 days old, **C**), met4 (#355, 201 days old, **D**), met5 (#1162, 232 days old, **E**), met6 (#1465, 253 days old, **F**), met7 (#354, 292 days old, **G**). H&E (left image), AR (middle image) and pan-cytokeratin (right image). Scale bars are 200µm (apart from **F**, peritoneum, 500µm).
