## Supplementary Figure 3 for "Plexin-B1 mutation drives prostate cancer metastasis"

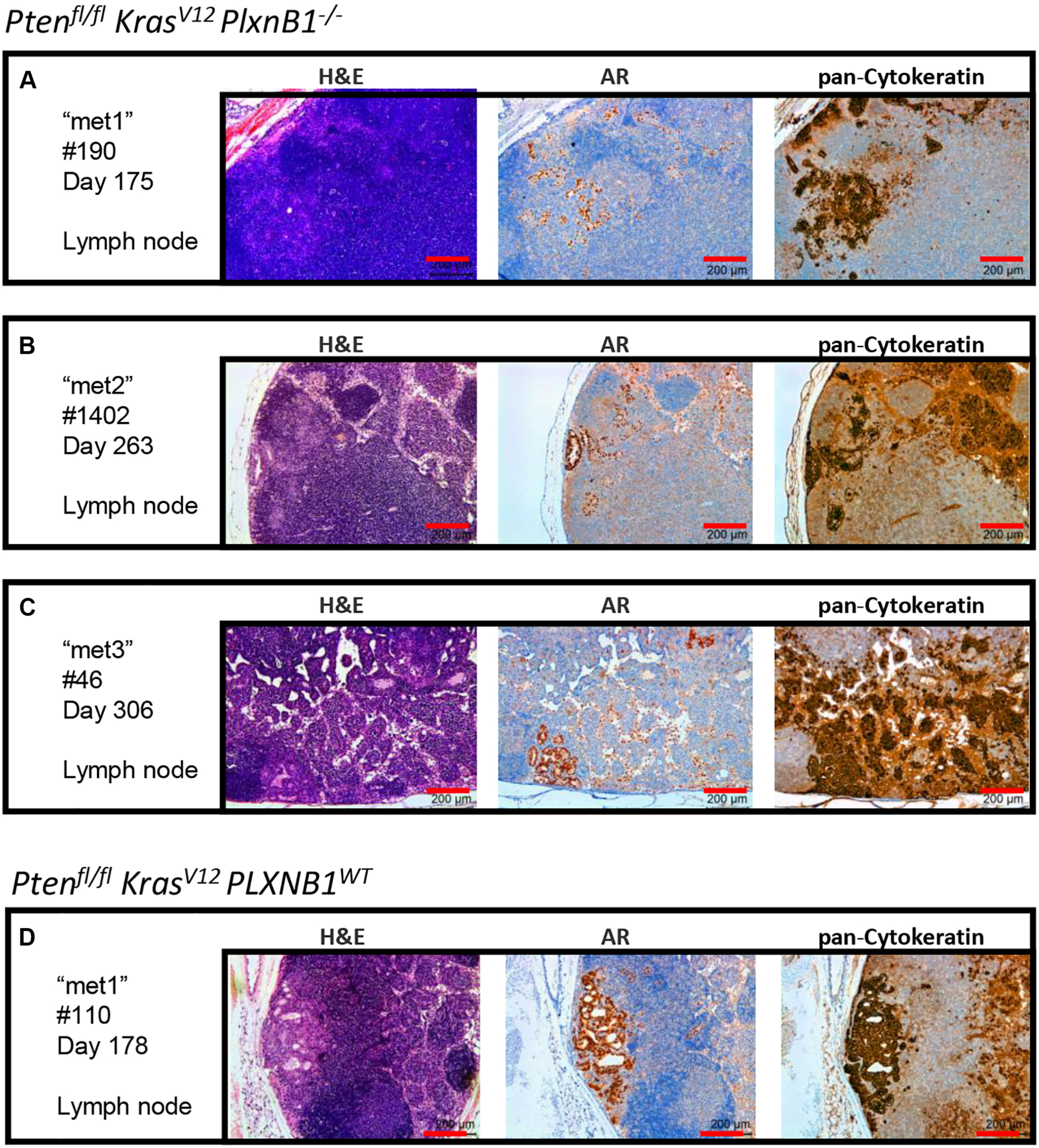
Pten fl/fl Kras V12 PlxnB1 -/-
A
"met1"
#190
Day 175
Lymph node
H&E
AR
pan-Cytokeratin
200 µm
B
"met2"
#1402
Day 263
Lymph node
C
"met3"
#46
Day 306
Lymph node
Pten fl/fl Kras V12 PLXNB1 WT
D
"met1"
#110
Day 178
Lymph node

**Supplementary Figure 3**. **Metastatic deposits in *Pten*$^{fl/fl}$*Kras*$^{G12V}$*PlxnB1*$^{-/-}$ mouse cohort and single *Pten*$^{fl/fl}$*Kras*$^{G12V}$*PLXNB1*$^{WT}$ mouse stained for H&E, androgen receptor (AR) and pan-cytokeratin. (A-C)** Metastatic deposits were observed in 3 *Pten*$^{fl/fl}$ *Kras*$^{V12}$ *PlxnB1*$^{-/-}$ cohort animals, met1 (#190, 175 days old, **A**), met2 (#1402, 263 days old, **B**), met3 (#46, 306 days old, **C**). H&E (left image), AR (middle image) and pan-cytokeratin (right image). Scale bars are 200µm. **(D)** Metastatic deposits were observed in a single *Pten*$^{fl/fl}$ *Kras*$^{V12}$ *PLXNB1*$^{WT}$ cohort animal, met1 (#110, 178 days old). Sections were stained for H&E (left image), AR (middle image) and pan-cytokeratin (right image). Scale bars are 200µm.
