## Supplementary Figure 4 for "Plexin-B1 mutation drives prostate cancer metastasis"

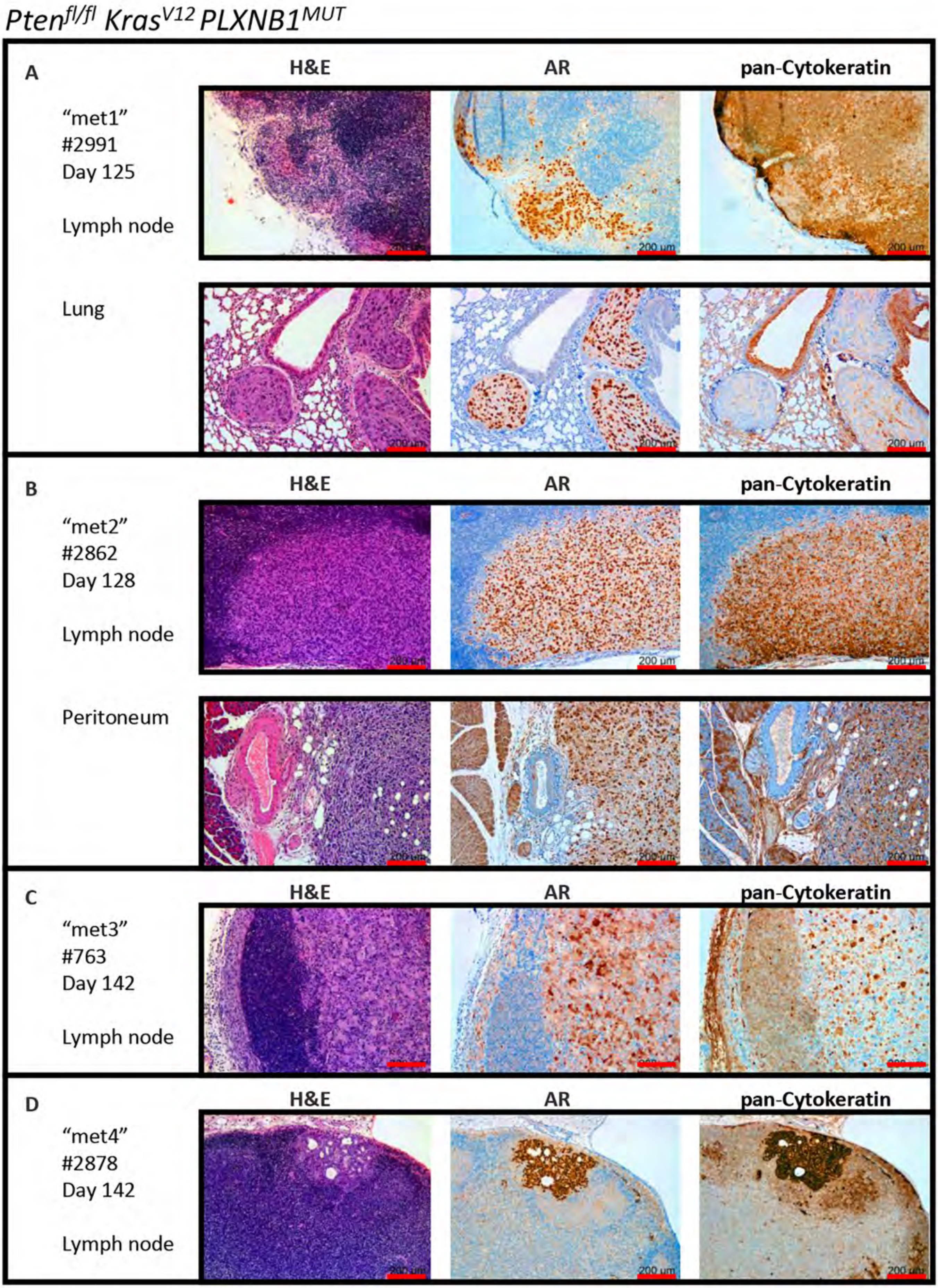
$Pten^{fl/fl}$ $Kras^{V12}$ $PLXNB1^{MUT}$
A
H&E
AR
pan-Cytokeratin
"met1"
#2991
Day 125
Lymph node
Lung
200 µm
B
H&E
AR
pan-Cytokeratin
"met2"
#2862
Day 128
Lymph node
Peritoneum
C
H&E
AR
pan-Cytokeratin
"met3"
#763
Day 142
Lymph node
D
H&E
AR
pan-Cytokeratin
"met4"
#2878
Day 142
Lymph node

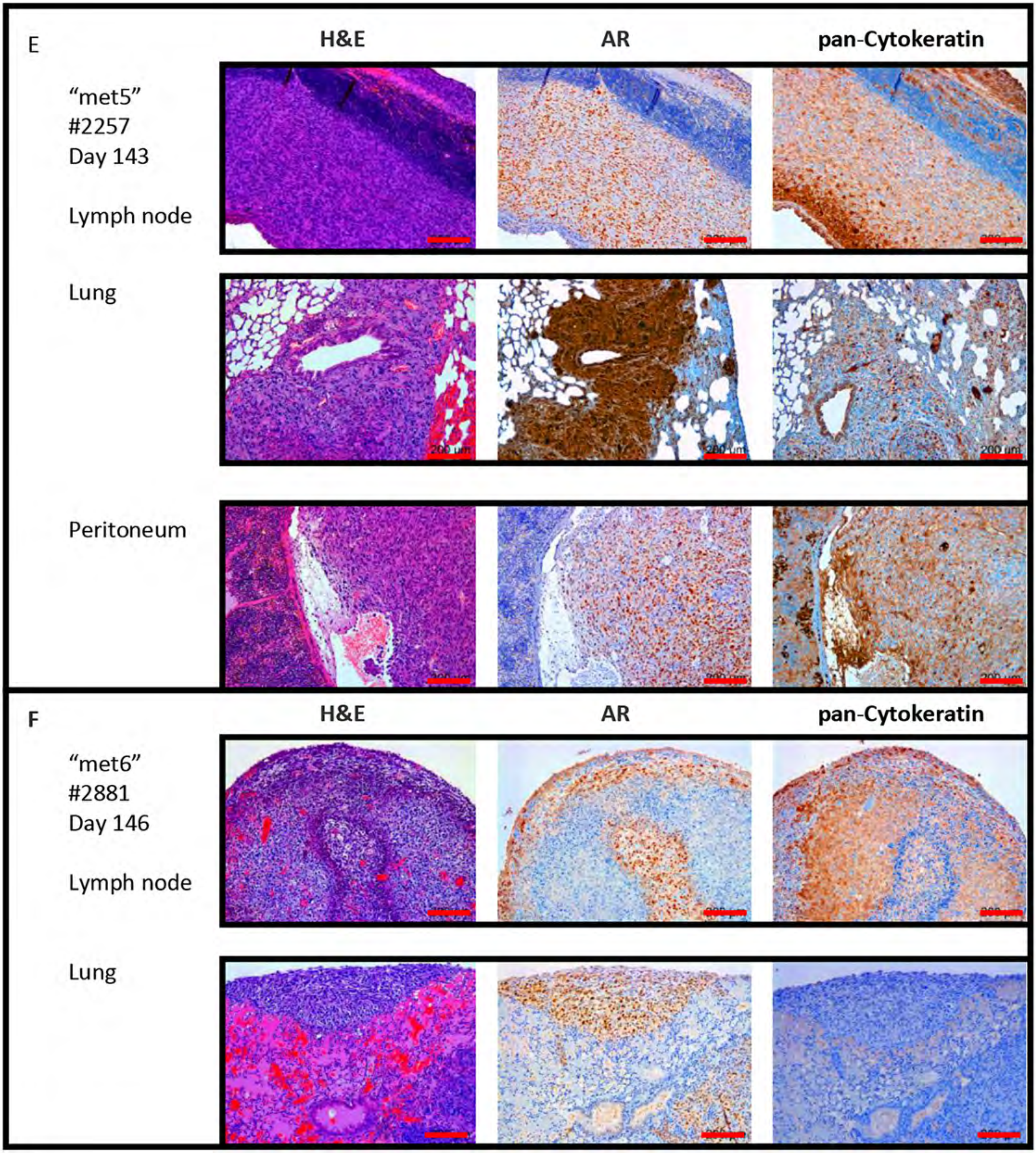
E
H&E
AR
pan-Cytokeratin
"met5"
#2257
Day 143
Lymph node
Lung
Peritoneum
F
H&E
AR
pan-Cytokeratin
"met6"
#2881
Day 146
Lymph node
Lung

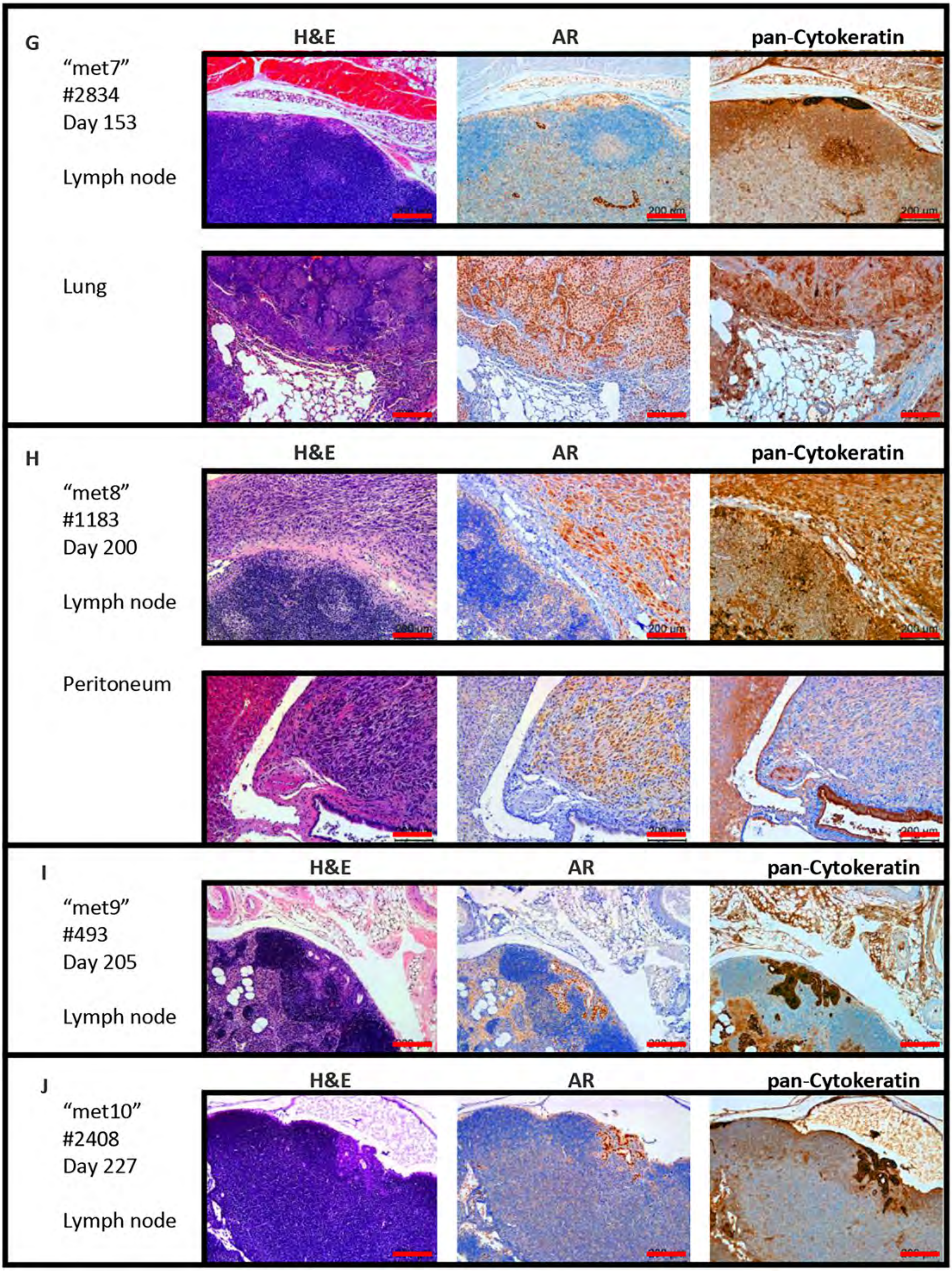
G
H&E
AR
pan-Cytokeratin
"met7"
#2834
Day 153
Lymph node
Lung
H
H&E
AR
pan-Cytokeratin
"met8"
#1183
Day 200
Lymph node
Peritoneum
I
H&E
AR
pan-Cytokeratin
"met9"
#493
Day 205
Lymph node
J
H&E
AR
pan-Cytokeratin
"met10"
#2408
Day 227
Lymph node

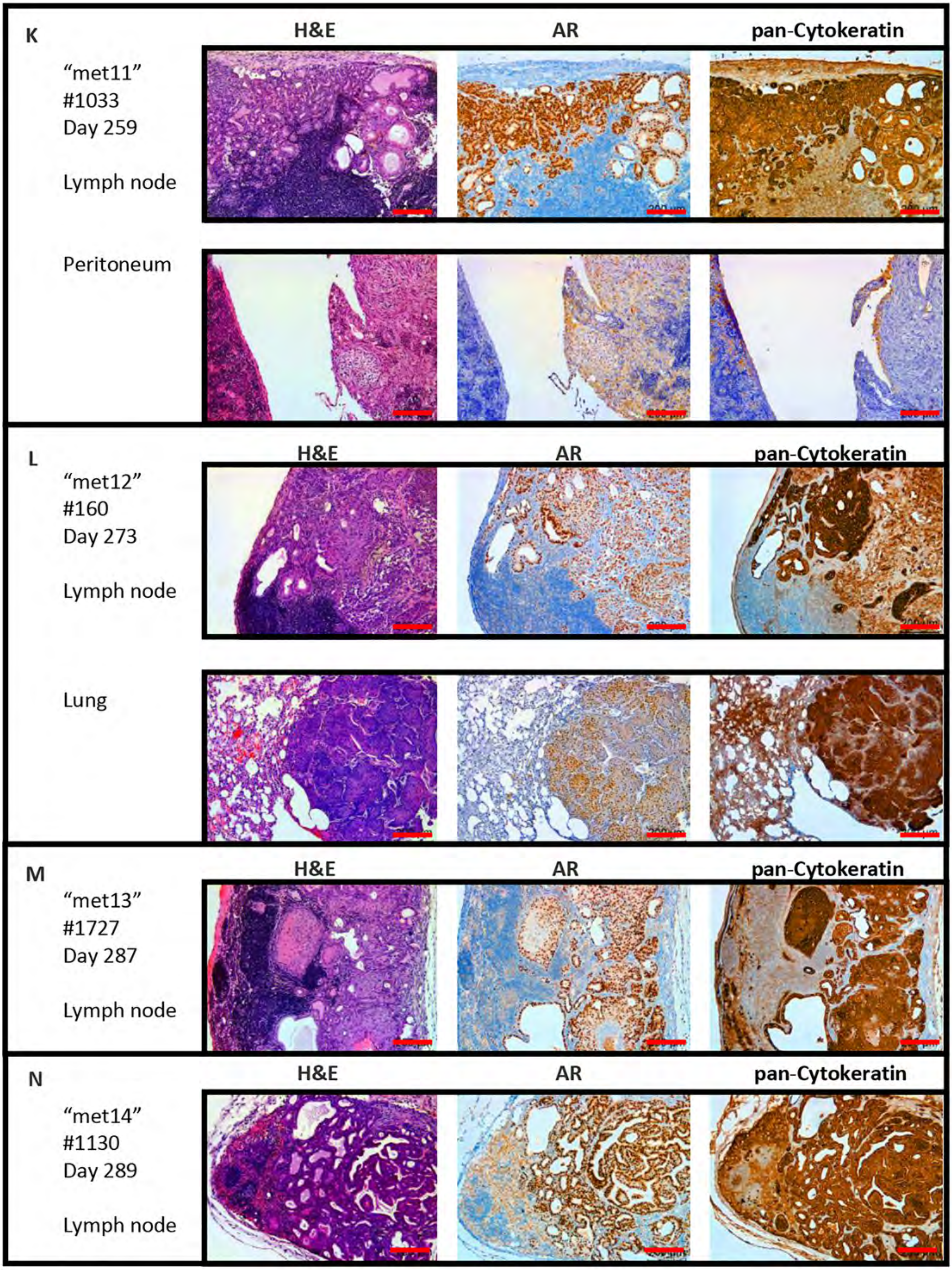
K
H&E
AR
pan-Cytokeratin
“met11”
#1033
Day 259
Lymph node
Peritoneum
L
H&E
AR
pan-Cytokeratin
“met12”
#160
Day 273
Lymph node
Lung
M
H&E
AR
pan-Cytokeratin
“met13”
#1727
Day 287
Lymph node
N
H&E
AR
pan-Cytokeratin
“met14”
#1130
Day 289
Lymph node

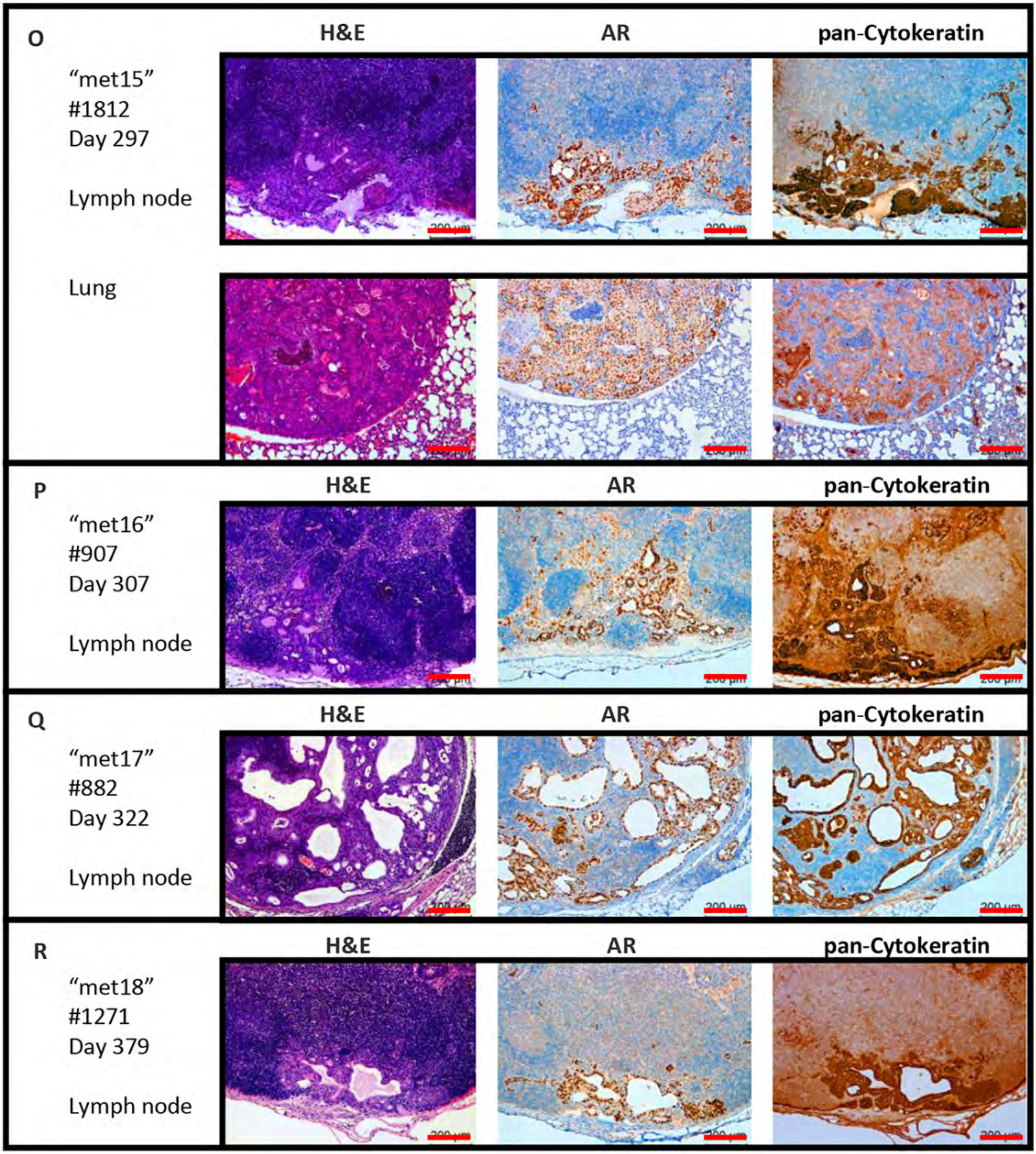
O
H&E
AR
pan-Cytokeratin
"met15"
#1812
Day 297
Lymph node
Lung
P
H&E
AR
pan-Cytokeratin
"met16"
#907
Day 307
Lymph node
Q
H&E
AR
pan-Cytokeratin
"met17"
#882
Day 322
Lymph node
R
H&E
AR
pan-Cytokeratin
"met18"
#1271
Day 379
Lymph node

**Supplementary Figure 4**. **Metastatic deposits in *Pten*$^{fl/fl}$*Kras*$^{G12V}$*PLXNB1*$^{MUT}$ mouse cohort stained for H&E, androgen receptor (AR) and pan-cytokeratin.** Metastatic deposits were observed in 18 *Pten*$^{fl/fl}$ *Kras*$^{V12}$ *PLXNB1*$^{MUT}$ cohort animals, met1 (#2991, 125 days old, **A**), met2 (#2862, 128 days old, **B**), met3 (#763, 142 days old, **C**), met4 (#2878, 142 days old, **D**), met5 (#2257, 143 days old, **E**), met6 (#2881, 146 days old, **F**), met7 (xplr2834, 153 days old, **G**), met8 (#1183, 200 days old,**H**), met9 (#493, 205 days old,**I**), met10 (#2408, 227 days old,**J**), met11 (#1033, 259 days old, **K**), met12 (#160, 273 days old, **L**), met13 (#1727, 287 days old, **M**), met14 (#1130, 289 days old, **N**), met15 (#1812, 297 days old, **O**), met16 (#907, 307 days old, **P**), met 17 (#882, 322 days old, **Q**), met18 (#1271, 379 days old, **R**). H&E (left image), AR (middle image) and pan-cytokeratin (right image). Scale bars are 200µm.
