## Supplementary Figure 5 for "Plexin-B1 mutation drives prostate cancer metastasis"

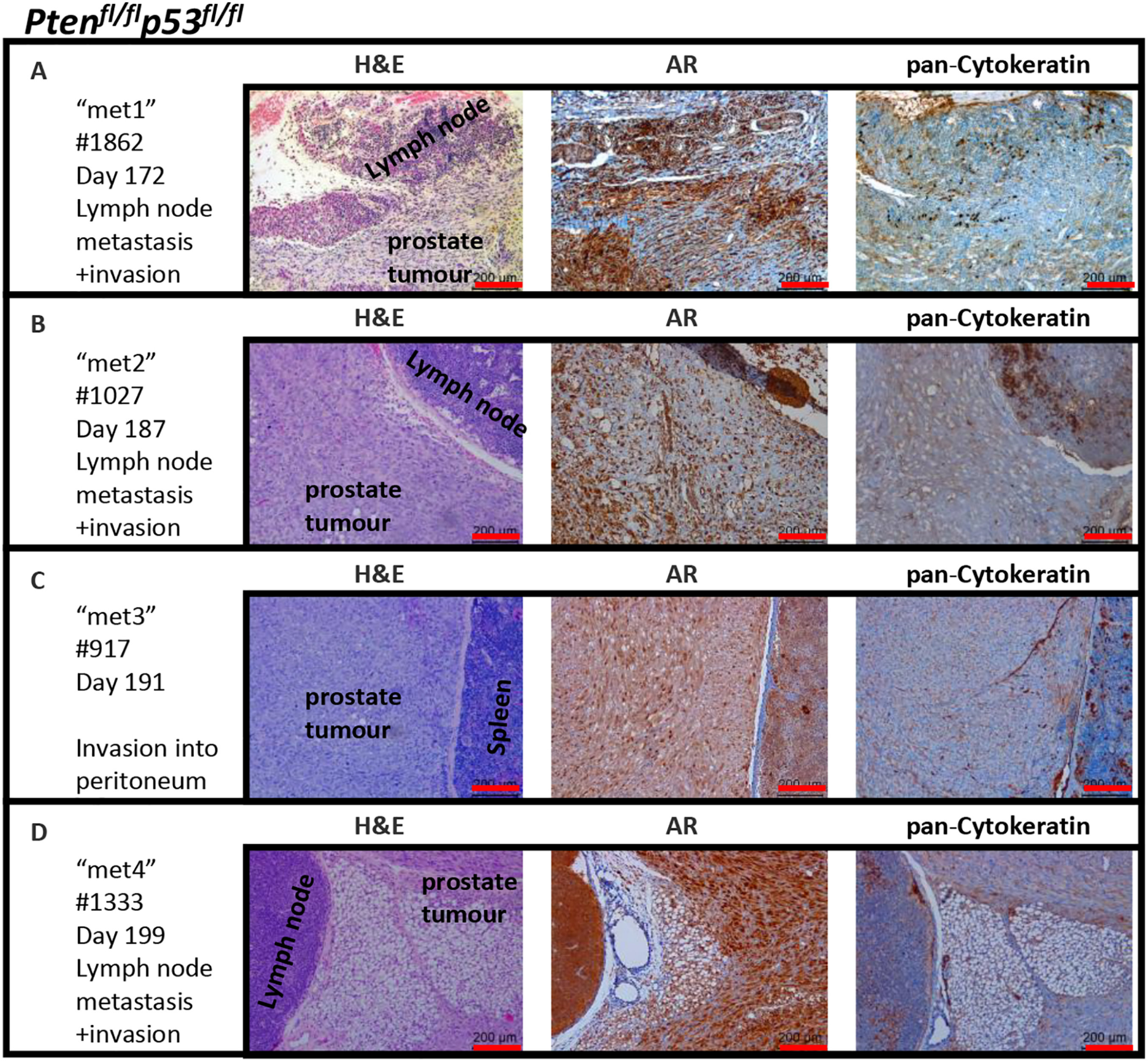

Pten$^{fl/fl}$p53$^{fl/fl}$
A
"met1"
#1862
Day 172
Lymph node metastasis +invasion
H&E
AR
pan-Cytokeratin
Lymph node
prostate tumour
200 µm
B
"met2"
#1027
Day 187
Lymph node metastasis +invasion
H&E
AR
pan-Cytokeratin
Lymph node
prostate tumour
200 µm
C
"met3"
#917
Day 191
Invasion into peritoneum
H&E
AR
pan-Cytokeratin
prostate tumour
Spleen
200 µm
D
"met4"
#1333
Day 199
Lymph node metastasis +invasion
H&E
AR
pan-Cytokeratin
Lymph node
prostate tumour
200 µm

**Supplementary Figure 5**. **Metastatic deposits in *Pten*$^{fl/fl}$*p53*$^{fl/fl}$ mouse cohort stained for H&E, androgen receptor (AR) and pan-cytokeratin.** Metastatic deposits were observed in four *Pten*$^{fl/fl}$*p53*$^{fl/fl}$ cohort animals, met1 (#1862, 172 days old, **A** ), met2 (#1027, 187 days old, **B**), met3 (#917, 191 days old, **C**), met4 (#1333, 199 days old, **D**). H&E (left image), AR (middle image) and pan-cytokeratin (right image). Scale bars are 200µm.
