## Supplementary Figure 6 for "Plexin-B1 mutation drives prostate cancer metastasis"

*$Pten^{fl/fl}p53^{fl/fl}$ $PLXNB1^{MUT}$*

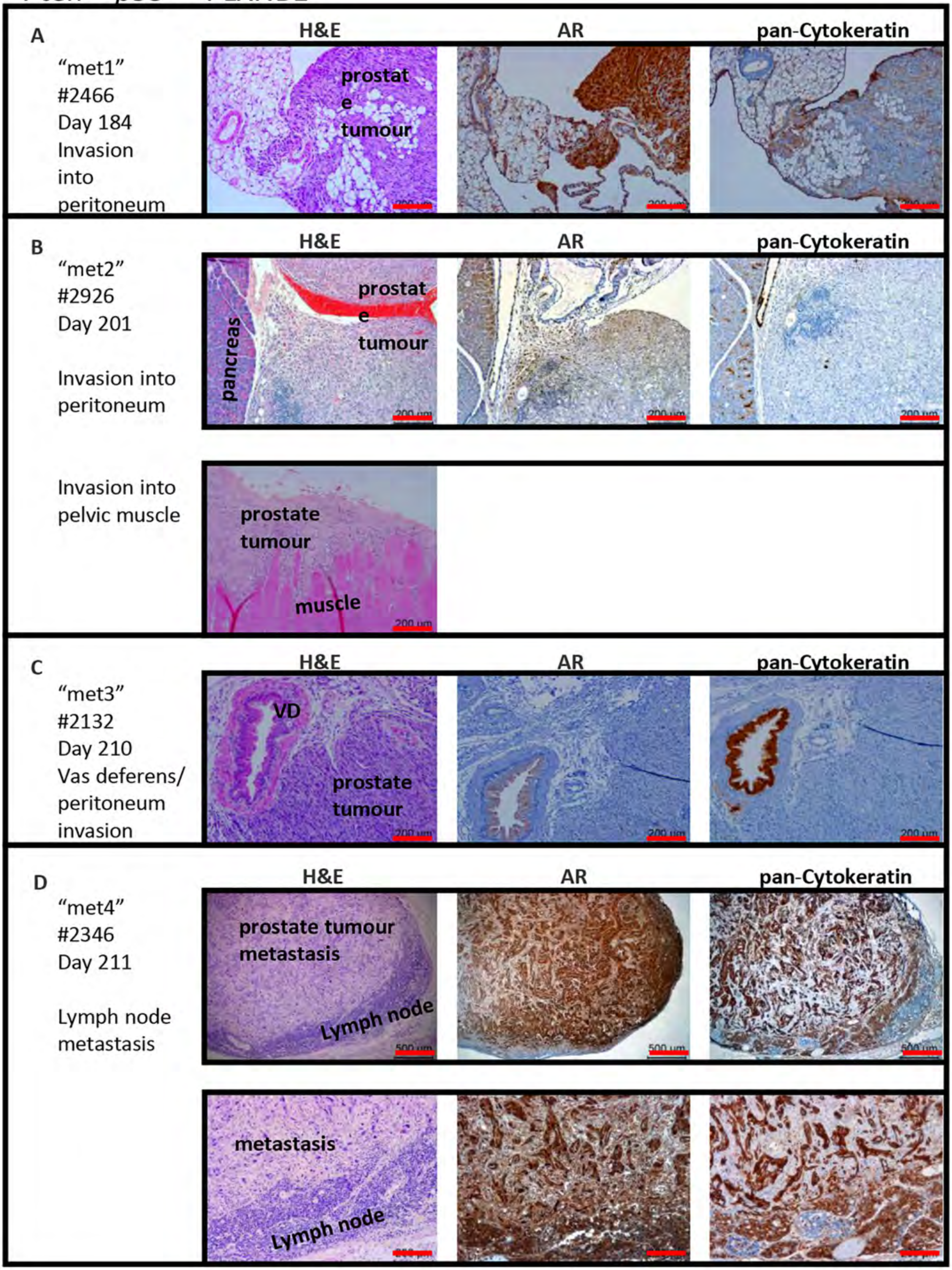

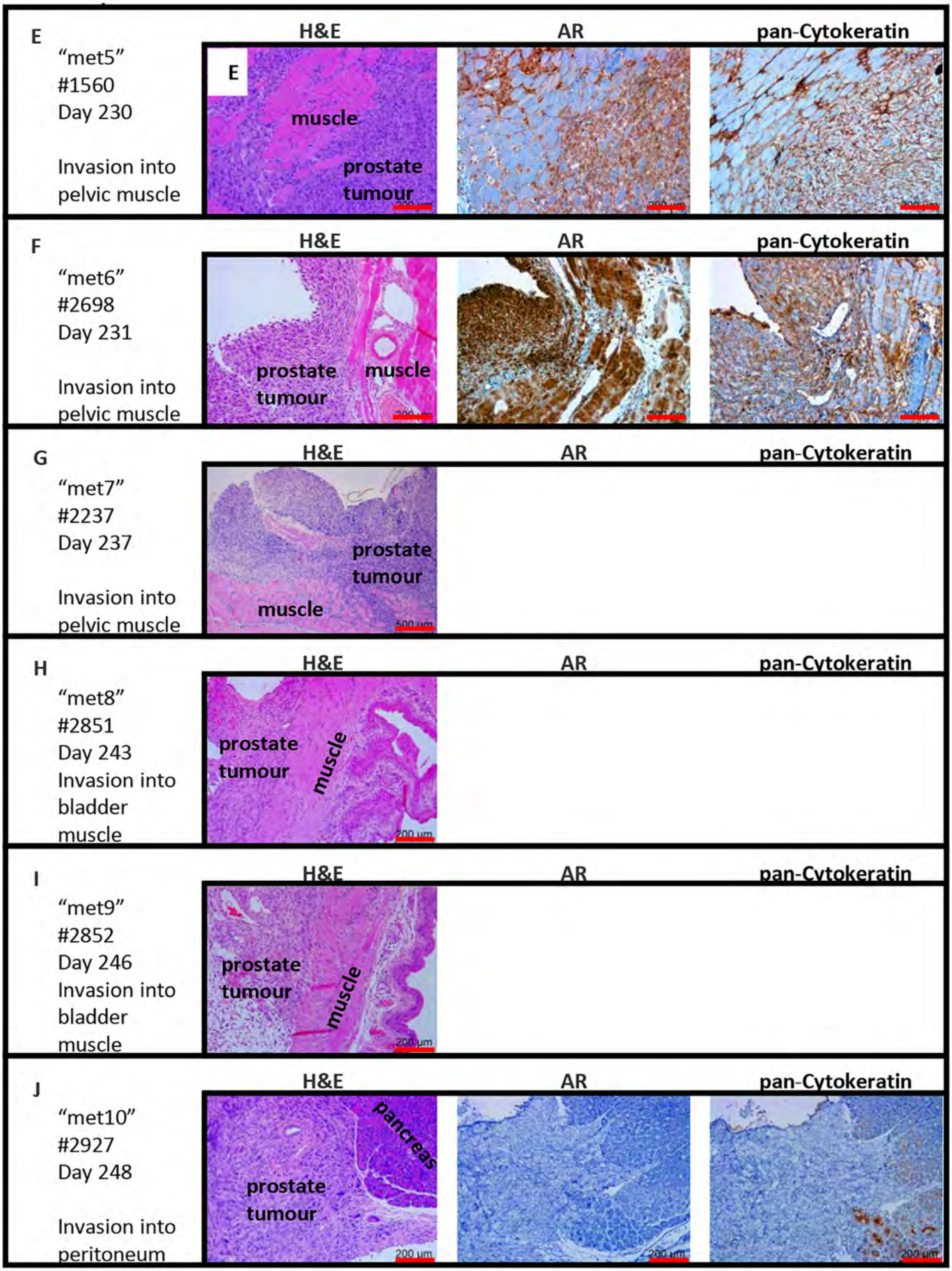
E
"met5"
#1560
Day 230
Invasion into pelvic muscle
H&E
AR
pan-Cytokeratin
E
muscle
prostate tumour
F
"met6"
#2698
Day 231
Invasion into pelvic muscle
H&E
AR
pan-Cytokeratin
prostate tumour
muscle
G
"met7"
#2237
Day 237
Invasion into pelvic muscle
H&E
AR
pan-Cytokeratin
prostate tumour
muscle
500 µm
H
"met8"
#2851
Day 243
Invasion into bladder muscle
H&E
AR
pan-Cytokeratin
prostate tumour
muscle
200 µm
I
"met9"
#2852
Day 246
Invasion into bladder muscle
H&E
AR
pan-Cytokeratin
prostate tumour
muscle
200 µm
J
"met10"
#2927
Day 248
Invasion into peritoneum
H&E
AR
pan-Cytokeratin
pancreas
prostate tumour
200 µm
200 µm
200 µm

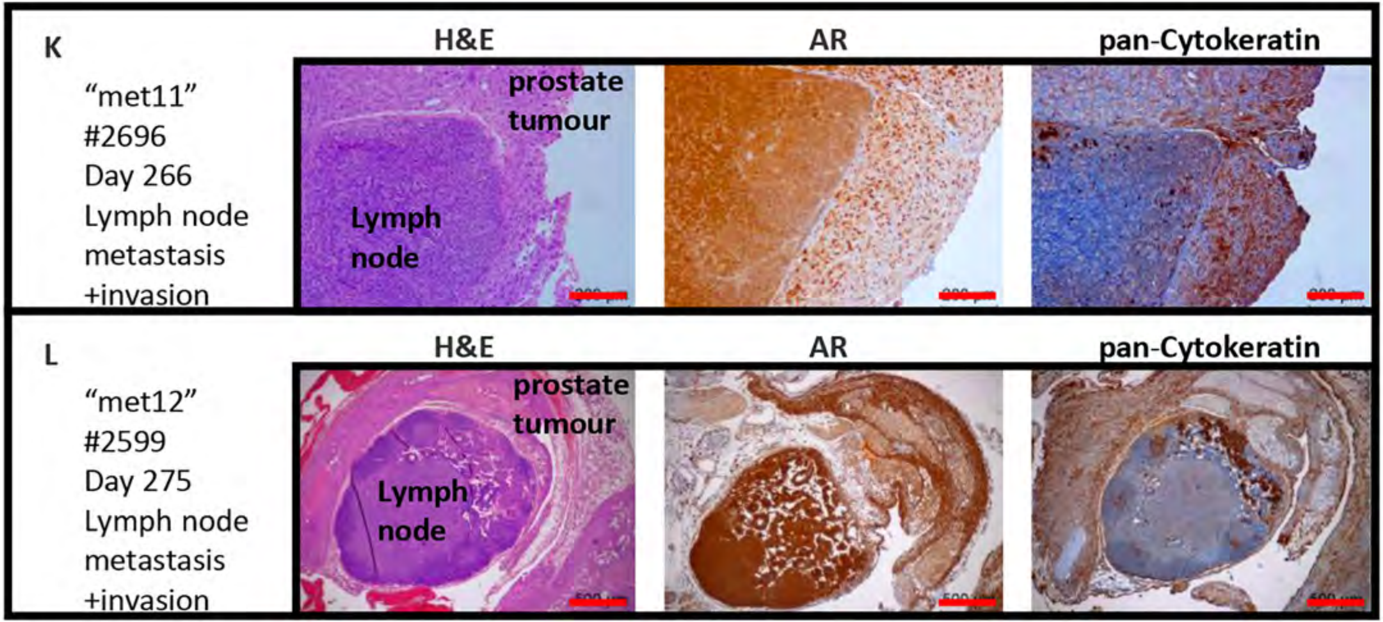
K
"met11"
#2696
Day 266
Lymph node
metastasis
+invasion
H&E
AR
pan-Cytokeratin
prostate
tumour
Lymph
node
L
"met12"
#2599
Day 275
Lymph node
metastasis
+invasion
H&E
AR
pan-Cytokeratin
prostate
tumour
Lymph
node

**Supplementary Figure 6**. **Metastatic deposits in *Pten*$^{fl/fl}$*p53*$^{fl/fl}$*PLXNB1*$^{MUT}$ mouse cohort stained for H&E, androgen receptor (AR) and pan-cytokeratin.** Metastatic deposits were observed in twelve *Pten*$^{fl/fl}$*p53*$^{fl/fl}$ *PLXNB1*$^{MUT}$ cohort animals, met1 (#2466, 172 days old, **A** ), met2 (#2926, 201 days old, **B**), met3 (#2132, 210 days old, **C**), met4 (#2346, 211 days old, **D**) met5 (#1560, 230 days old, **E**) met6 (#2698, 231 days old, **F**) met7 (#2237, 237 days old, **G**) met8 (#2851, 243 days old, **H**) met9 (#2852, 246 days old, **I**) met10 (#2927, 248 days old, **J**) met11 (#2696, 266 days old, **K**) met12 (#2599, 275 days old, **L**). H&E (left image), AR (middle image) and pan-cytokeratin (right image). Scale bars are 200µm apart from **D** (upper row), **G** and **L** (500 µm).
