## Supplementary Figure 7 for "Plexin-B1 mutation drives prostate cancer metastasis"

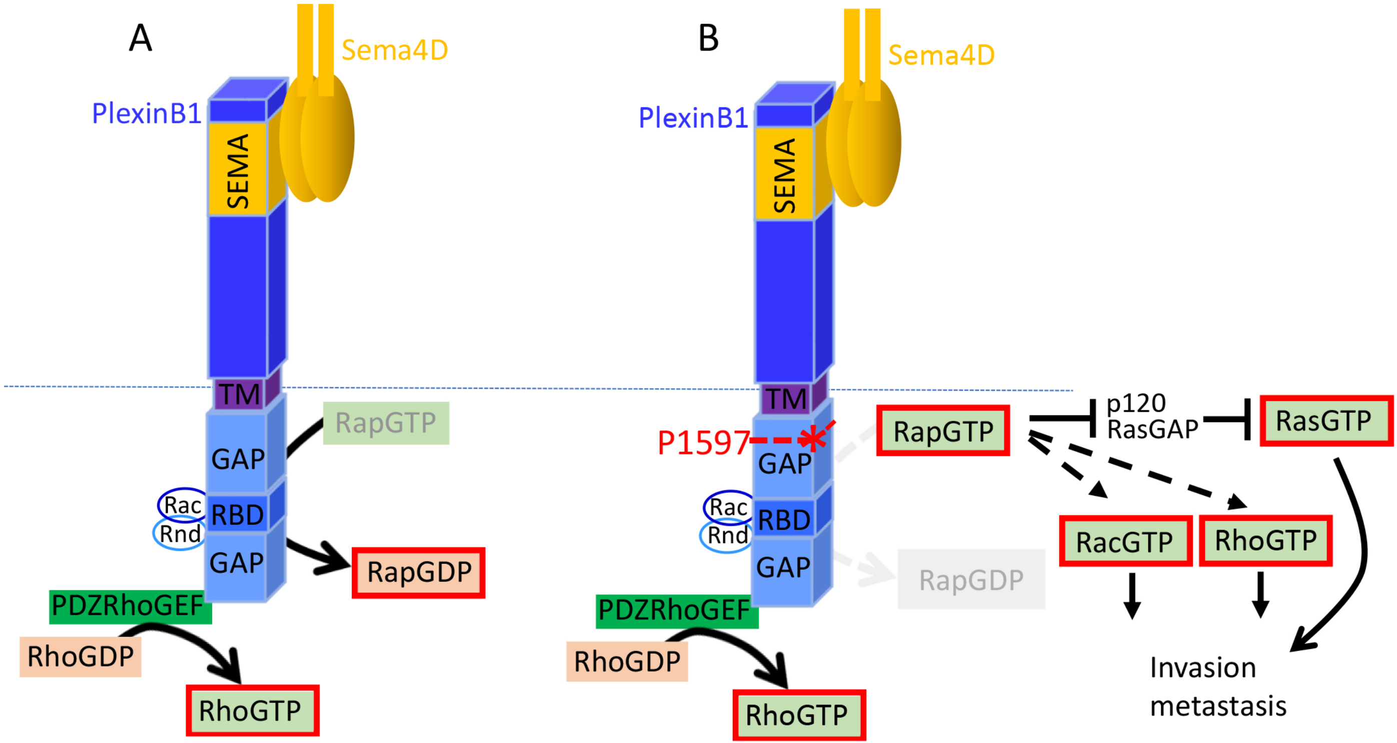
A
Sema4D
PlexinB1
SEMA
TM
RapGTP
GAP
Rac
RBD
Rnd
GAP
RapGDP
PDZRhoGEF
RhoGDP
RhoGTP
B
Sema4D
PlexinB1
SEMA
TM
P1597
RapGTP
p120
RasGAP
RasGTP
GAP
Rac
RBD
Rnd
GAP
RacGTP
RhoGTP
RapGDP
PDZRhoGEF
RhoGDP
RhoGTP
Invasion
metastasis

**Supplementary Figure 7. Model of effect of P1597L mutation on PlexinB1 signalling**. (**A)** Wild type PlexinB1 regulates several small GTPases including Rap. GTP-bound Rap1 represses p120RasGAP leading to Ras activation (42); Rap1 also activates Rac and Rho signalling promoting tumour growth, cell motility and invasion. Wild-type PlexinB1 acts as a GTPase activating protein (GAP) for Rap, catalysing the conversion of RapGTP to inactive RapGDP. WT PlexinB1, through its activity as a RapGAP, thereby reduces Rap activation, decreasing Ras, Rac, and Rho activity and suppressing cell motility, invasion and MLC2 phosphorylation. **(B)** The P1597L mutation in the GAP domain of PlexinB1 is predicted to disrupt the GAP activity of PlexinB1. Consequently, RapGTP is not inactivated by mutant PlexinB1, allowing Rap in its active GTP bound form to promote Ras, Rho and Rac activation leading to MLC2 phosphorylation, invasion and metastasis. TM, transmembrane domain; SEMA, SEMA domain; GAP, GTPase activating protein domain; RBD, Rho Binding domain; *position of P1597L mutation.
