## Supplementary Table 1 for "Plexin-B1 mutation drives prostate cancer metastasis"

| Study | Sample analysis method | Percentage of patients with altered Plexin B1 expression | | Total number of patient mRNA expression profiles |
|---|---|---|---|---|
| | | z-score=±2 | | |
| | | high | low | |
| Prostate Adenocarcinoma (MSKCC, Cancer Cell 2010) doi:10.1016/j.ccr.2010.05.026. | mRNA z-scores compared to expression in normal prostate samples | 7.33 | 22.67 | 150 |
| | mRNA z-scores (metastatic tumors) compared to expression in normal prostate samples | 15.79 | 26.32 | 19 |
| | mRNA z-scores (metastatic tumors, non-castrate) compared to expression in normal prostate samples | 12.50 | 12.50 | 8 |
| | mRNA z-scores (metastatic tumors, castrate) compared to expression in normal prostate samples | 18.18 | 36.36 | 11 |
| Neuroendocrine Prostate Cancer (Multi-Institute, Nat Med 2016) doi:10.1038/nm.4045. | mRNA expression z-scores relative to diploid samples | 5.71 | 0.00 | 35 |
| | mRNA expression z-scores relative to all samples (log microarray) | 0.00 | 45.71 | 35 |
| Prostate Adenocarcinoma (Fred Hutchinson CRC, Nat Med 2016) doi: 10.1038/nm.4053 | mRNA expression z-scores relative to diploid samples | 12.70 | 7.94 | 63 |
| | mRNA expression z-scores relative to all samples (log microarray) | 6.35 | 6.35 | 63 |
| Prostate Adenocarcinoma (SMMU, Eur Urol 2017) doi:10.1016/j.eururo.2017.08.027. | mRNA Expression z-scores relative to diploid samples (RNA Seq FPKM) | 4.62 | 0.00 | 65 |
| | mRNA expression z-scores relative to all samples (log RNA Seq FPKM) | 1.54 | 3.08 | 65 |
| Prostate Cancer (DKFZ, Cancer Cell 2018) doi: 10.1016/j.ccell.2018.10.016. | mRNA expression z-scores relative to all samples (log RNA Seq RPKM) | 4.21 | 2.11 | 95 |
| Metastatic Prostate Adenocarcinoma (SU2C/PCF Dream Team, PNAS 2019) doi: 10.1073/pnas.1902651116 | mRNA expression z-scores relative to diploid samples (FPKM capture) | 4.88 | 0.00 | 205 |
| | mRNA expression z-scores relative to all samples (log FPKM capture) | 0.98 | 1.46 | 205 |
| Prostate Adenocarcinoma (TCGA, PanCancer Atlas) doi:10.1016/j.cell.2015.10.025 | mRNA expression z-scores relative to diploid samples (RNA Seq V2 RSEM) | 2.43 | 0.41 | 493 |
| | mRNA expression z-scores relative to all samples (log RNA Seq V2 RSEM) | 0.61 | 3.85 | 493 |

**Supplementary Table 1. Plexin-B1 expression data in human prostate cancer.** The table shows the percentage of patients with altered (high or low) Plexin-B1 expression in 7 prostate cancer genomics projects of gene expression (RNAseq and microarray) currently available through cBioPortal(48,49). A z-score=±2 threshold (2 standard deviations above or below the mean) was used to calculate the percentage.
